# Denitrification kinetics indicates nitrous oxide uptake is unaffected by electron competition in Accumulibacter

**DOI:** 10.1101/2020.05.14.092429

**Authors:** Roy Samarpita, Pradhan Nirakar, NG How Yong, Wuertz Stefan

**Author notes:** Corresponding author: Stefan Wuertz, Singapore Centre for Environmental Life, Sciences Engineering, Nanyang Technological University, Singapore.

## Abstract

Denitrifying phosphorus removal is a cost and energy efficient treatment technology that relies on polyphosphate accumulating organisms (DPAOs) utilizing nitrate or nitrite as terminal electron acceptor. Denitrification is a multistep process and many organisms do not possess the complete pathway, leading to the accumulation of intermediates such as nitrous oxide (N_2_O), a potent greenhouse gas and ozone depleting substance. *Candidatus* Accumulibacter organisms are prevalent in denitrifying phosphorus removal processes and, according to genomic analyses, appear to vary in their denitrification abilities based on their lineage. Yet, denitrification kinetics and nitrous oxide accumulation by Accumulibacter after long-term exposure to either nitrate or nitrite as electron acceptor have never been compared. We investigated the preferential use of the nitrogen oxides involved in denitrification and nitrous oxide accumulation in two enrichments of Accumulibacter and a competitor – the glycogen accumulating organism *Candidatus* Competibacter. A metabolic model was modified to predict phosphorus removal and denitrification rates when nitrate, nitrite or N_2_O were added as electron acceptors in different combinations. Unlike previous studies, no N_2_O accumulation was observed for Accumulibacter in the presence of multiple electron acceptors. Electron competition did not affect denitrification kinetics or N_2_O accumulation in Accumulibacter or Competibacter. Despite the presence of sufficient internal storage polymers (polyhydroxyalkanoates, or PHA) as energy source for each denitrification step, the extent of denitrification observed was dependent on the dominant organism in the enrichment. Accumulibacter showed complete denitrification and N_2_O utilization, whereas for Competibacter denitrification was limited to reduction of nitrate to nitrite. These findings indicate that DPAOs can contribute to lowering N_2_O emissions in the presence of multiple electron acceptors under partial nitritation conditions.

## 1. INTRODUCTION

Biological nutrient removal processes are regarded as cost-effective, sustainable solutions to tackle excess carbon, phosphorus (P) and nitrogen (N) in wastewater. However, complete P and N removal usually requires high aeration, accounting for more than 60% of the overall power consumption at a wastewater treatment plant (WWTP) [1]. To meet rigorous effluent discharge limits, external carbon dosage is often required to achieve complete denitrification resulting in increased operational costs [2]. In this regard, denitrifying phosphorus removal (DPR) has been identified as an optimal treatment solution due to its reduced energy consumption, sludge wastage and carbon requirement for nutrient removal.

In the DPR process, sludge is cycled through anaerobic-anoxic conditions allowing denitrifying phosphorus accumulating organisms (DPAOs) such as *Candidatus* Accumulibacter phosphatis (hereafter referred to as Accumulibacter) to take up organic carbon under anaerobic conditions by utilizing stored polyphosphate as a source of energy. In the subsequent anoxic phase, electron acceptors such as nitrate or nitrite are reduced to meet growth and metabolic requirements [3-5]. These conditions also lead to the growth of denitrifying glycogen accumulating organisms (DGAOs), which compete with DPAOs for organic carbon under anaerobic conditions, but do not aid in phosphorus removal in the subsequent phase; hence recognized as competitors to DPAOs. *Candidatus* Competibacter phosphatis (hereafter referred to as Competibacter) is one such DGAO commonly found in laboratory- and full-scale studies [6-9].

In terrestrial and aquatic ecosystems, the conversion of nitrous oxide (N_2_O) to nitrogen gas (E°^′^ = +1.35 V at pH 7) by nitrous oxide reductase (Nos) is the only known biological N_2_O attenuation process in the biosphere [10, 11]. Genomic evidence revealed that many organisms possess only a subset of this denitrification pathway, sometimes lacking the nitrous oxide reductase gene (*nos*), resulting in the accumulation of N_2_O [11, 12]. This is particularly important in engineered systems such as WWTPs where denitrification is implemented to remove nitrogen from influent wastewater [13, 14]. To this end, a genomic comparison of the denitrification pathway in Accumulibacter clades IA, IIA, IIB, and IIC showed significant differences and suggested that some clades were incapable of reducing N_2_O [15-17]. Likewise, there is evidence that phylogenetically similar members of Competibacter have distinct denitrification capabilities [7, 18-20]. However, most metabolic models ignore the differences in denitrification capabilities of PAOs and GAOs and split the total nitrogen reduced between anoxic and aerobic phases [21-24]. Such an approach overlooks differences in phosphorus removal rates by different Accumulibacter clades that arise due to affinity towards certain nitrogen oxides. In addition to differences in denitrification abilities, environmental factors have been associated with increased N_2_O release from WWTPs [24-27]. For heterotrophic organisms, studies demonstrated that the presence of nitrogen oxides (i.e., nitrate or nitrite) coupled with free nitrous acid (FNA) accumulation had a negative effect on the nitrous oxide reduction potential [28]. Nitrous oxide accumulation is also affected by electron competition, which results in a higher flow of electrons to one step of the denitrification pathway, rather than electrons being distributed to all denitrification enzymes [25, 28, 29]. In the context of Accumulibacter, slow polyhydroxyalkanoates (PHA) degradation has often been reported as a reason for N_2_O release from DPR systems, although others have been unable to establish a link between the two [30, 31]. Thus, it is unclear whether N_2_O emission from the DPR process is due to environmental factors or an incomplete denitrification pathway in dominant denitrifying organisms.

Metabolic models have been widely used to predict nutrient removal in wastewater treatment [21-23, 32]. So far, models predicting N_2_O accumulation have been developed solely based on external nitrate dosage as terminal electron acceptor [18, 33-36]. However, various additional nitrogen oxides (NO_x_) occur simultaneously in biological wastewater treatment, and each can serve as terminal electron acceptor. This necessitates validation of metabolic models to evaluate electron competition and denitrification kinetics in organisms utilizing PHA. The preference for different NO_x_ has also been shown to affect oxidative phosphorylation and phosphorus transport across cell membranes [26, 36, 37]. Consequently, it is necessary to consider differences in the denitrification pathway and the effect of specific long-term enrichment conditions (nitrate or nitrite as electron acceptor) in modelling approaches.

In this study, we compared the denitrification kinetics of two Accumulibacter enrichments acclimated to different anoxic conditions. We tested the effect of electron competition on nitrous oxide accumulation in the presence of multiple electron acceptors. These results for Accumulibacter were contrasted with denitrification characteristics of a Competibacter enrichment. The objectives were to (i) identify any accumulation of intermediates in the denitrification process that would signify a preference for certain terminal electron acceptors, (ii) compare the role of electron competition in nitrous oxide accumulation, and (iii) validate an existing metabolic model to predict the mechanism of phosphorus uptake and nitrogen oxide accumulation in the presence of multiple electron acceptors. We investigated three electron acceptors (nitrate, nitrite and nitrous oxide) dosed in seven different combinations under similar test conditions for all enrichments. We hypothesized that the preference for certain NO_x_ compounds would be driven either by electron competition or the extent of denitrification performed by the dominant organism in each enrichment culture.

## 2. MATERIALS AND METHODS

### 2. 1. Reactor operation for enrichment cultures

We operated three lab-scale aerobic granular sludge sequencing batch reactors (SBRs), each with a 2-L working volume and 200 mL headspace, to obtain Accumulibacter and Competibacter enrichments. Accumulibacter was adapted to nitrate or nitrite as terminal electron acceptor under anoxic conditions leading to two different enrichments, Accumulibacter_nitrate_ and Accumulibacter_nitrite_, respectively. The third reactor enriched Competibacter fed with nitrate under anoxic conditions. The reactors were maintained at a constant temperature of 30°C using a heating jacket, and a cycle length of 6 h consisting of 30 min feeding, 60 min anaerobic react, 120 min anoxic react, 120 min aerobic react, 10 min settling and 20 min effluent discharge phases. During the anaerobic phase, 1 L of synthetic feed containing acetate and propionate (3:1 ratio on the basis of total influent chemical oxygen demand, or COD) as carbon source was pumped into each reactor at a rate of 33.33 mL/min.

Nitrogen gas was sparged from the bottom of the reactor at 2.0 L/min to provide sufficient mixing during the reaction phases and maintain anaerobic conditions. Subsequently, in the anoxic phase nitrite or nitrate were fed in pulses. Nitrite and nitrate dosages were carefully adjusted to avoid their accumulation in the anoxic phase because this could affect microbial activity in the reactors. During the effluent discharge phase, 1 L of effluent was pumped out of the reactors, leading to a hydraulic retention time of 12 h and a volumetric exchange ratio of 50%. For each cycle, the pH was controlled at 7.5 to 8.0 with 0.5 M NaOH or 0.5 M HCl solution, and the dissolved oxygen (DO) concentration in the aerobic phase was varied from 1 to 1.5 mg O_2_/L. A programmable logic controller was linked to a SCADA interface for data visualization and storage. The bioreactors were assumed to be at pseudo-steady state at the start of the batch tests as no significant changes in mixed liquor volatile suspended solids (MLSS), volatile suspended solids (VSS) and phosphorus concentrations were observed for three consecutive solids retention times (SRTs). Cycle studies were performed regularly and samples were taken during each phase to monitor reactor performance. The operating conditions for the Accumulibacter_nitrite_, Accumulibacter_nitrate_ and Competibacter enrichment reactors were as follows:

#### Case 1: Accumulibacter_nitrite_

A lab-scale aerobic granular sludge SBR operated at 30°C and fed with synthetic wastewater in which the influent carbon source was a mixture of acetate and propionate (3:1 ratio for acetate and propionate on the basis of total influent COD) was used to enrich Accumulibacter. The synthetic wastewater consisted of 0.15 L solution A and 0.85 L solution B. Solution A (per litre) contained 3.5 g NaAc.3H_2_O and 0.7 mL of 99.5% propionic acid as the carbon source, 1.20 g MgSO_4_.7H_2_O, 0.19 g CaCl_2_.2H_2_O, 1.02 g NH_4_Cl, 0.01 g peptone, 0.01 g yeast and 4 mL of trace metals solution. The trace metals solution was prepared as described in Smolders et al. (1994) and consisted (per litre) of 0.15 g H_3_BO_3_, 1.5 g FeCl_3_.6H_2_O, 0.18 g KI, 0.12 g MnCl_2_.4H_2_O, 0.15 g CoCl_2_.6H_2_O, 0.06 g Na_2_MoO_4_.2H_2_O, 0.03 g CuSO_4_.5H_2_O, 0.12 g ZnSO_4_.7H_2_O and 10 g Ethylenediamine tetra-acetic acid (EDTA). Solution B contained 132 mg K_2_HPO_4_/L and 103 mg KH_2_PO_4_/L. Nitrification was inhibited by the addition of 2.38 mg allyl-N thiourea (ATU) per litre feed. After feeding, the mixed liquor contained 12 mg NH_4_-N/L, 300 mg COD/L and 15 mg PO_4_-P/L resulting in a COD to P ratio of 20:1. The anoxic phase was initiated by adding NaNO_2_ solution in pulses such that the total concentration in the reactor after each pulse was 10 mg NO_2_-N/L. The SRT varied from 12 to 15 d to maintain an MLSS between 2 and 2.5 g/L.

#### Case 2: Accumulibacter_nitrate_

Accumulibacter was enriched at 30°C in a lab-scale aerobic granular sludge SBR using the same enrichment protocol and synthetic wastewater as mentioned in Case I, Accumulibacter_nitrite_. The only difference in operation of this reactor was that the anoxic phase was initiated by adding KNO_3_ solution in pulses such that the final concentration in the reactor after each pulse was 15 mg NO_3_-N/L.

#### Case 3: Competibacter

Enrichment of Competibacter was achieved in a lab-scale aerobic granular sludge SBR operated at 30°C with synthetic feed containing acetate as sole carbon source. The synthetic wastewater was similar to that outlined for Case I (Accumulibacter_nitrite_) with the difference that solution A contained 8.5 g NaAc.3H_2_O and solution B contained trace amounts of K_2_HPO_4_ and KH_2_PO_4_, such that the resulting phosphorus concentration in the reactor after feeding was 2 mg P/L. After feeding, the mixed liquor contained 300 mg COD/L, 12 mg NH_4_-N/L and 2 mg PO_4_-P/L resulting in a COD:P of 150:1. The reactor operation phases were similar to those of the other reactors and KNO_3_ solution was introduced as a pulse into the reactor during the anoxic phase, resulting in a final concentration in the reactor after each pulse of 15 mg NO_3_-N/L.

### 2. 2. Batch tests to measure denitrification kinetics and N_2_O accumulation

A total of seven batch tests were performed under different terminal electron acceptor conditions for all enrichments. At the start of each batch test, mixed liquor was withdrawn from the reactor after completion of the anaerobic phase and immediately transferred to batch reactors with nitrogen gas sparging for 5 min to remove trace oxygen. The batch reactors had a total volume of 1.1 L and a working volume of 1 L. The reactor headspace was minimized in order to reduce stripping of nitrous oxide. For all tests, pH was controlled at 8.0 ± 0.1 using 0.5M HCl or 0.5M NaOH for Accumulibacter enrichments and 0.2M HCl or 0.2M NaOH for the Competibacter enrichment. Nitrous oxide reduction can be severely affected by FNA concentrations as low as 0.7 µg HNO_2_-N/L [37]. The aforementioned pH set-point was maintained during the batch tests to reduce susceptibility of sludge to the inhibitory effects of FNA. Once the tests began, no nitrogen was sparged in the batch reactor to minimize N_2_O stripping, and the mixed liquor was stirred slowly at a rate of 30 rpm to avoid oxygen intrusion. Dissolved N_2_O concentrations were monitored with an online N_2_O microsensor (N_2_O -R, Unisense A/S, Denmark). Fresh stock of N_2_O solution prepared by sparging Milli-Q water with 100% N_2_O gas for 5 min at room temperature (24-25°C) was used to calibrate the N_2_O probe as per the product instruction manual. N_2_O reduced at the cathode of the microsensor produced a current that was converted into a signal by the Picoammeter, and these readings were collected every second.

Different combinations of terminal electron acceptors were added for batch tests with the enrichment cultures (Table 1). In each test, nitrogen oxides were added as a pulse such that their concentrations in each batch test reactor ranged from 12 to 15 mg NO_x_-N/L. Four control tests were carried out in duplicate to determine the loss of N_2_O due to stripping or other abiotic phenomena. The losses were averaged and deducted from all Picoammeter readings to obtain the true biological N_2_O uptake as shown in the supplementary information (Appendix 2). The tests were conducted for 30 min and mixed liquor samples for nutrient analysis were collected every 5 min.

**Table 1.** Electron acceptors added in different batch tests

| Batch test | Electron acceptor | Intermediate measured |
| --- | --- | --- |
| A | N<sub>2</sub>O | N<sub>2</sub>O |
| B | NO<sub>3</sub><sup>-</sup> | NO<sub>3</sub><sup>-</sup>, NO<sub>2</sub><sup>-</sup>, N<sub>2</sub>O |
| C | NO<sub>2</sub><sup>-</sup> | NO<sub>2</sub><sup>-</sup>, N<sub>2</sub>O |
| D | NO<sub>3</sub><sup>-</sup>, NO<sub>2</sub><sup>-</sup> | NO<sub>3</sub><sup>-</sup>, NO<sub>2</sub><sup>-</sup>, N<sub>2</sub>O |
| E | NO<sub>3</sub><sup>-</sup>, N<sub>2</sub>O | NO<sub>3</sub><sup>-</sup>, NO<sub>2</sub><sup>-</sup>, N<sub>2</sub>O |
| F | NO<sub>2</sub><sup>-</sup>, N<sub>2</sub>O | NO<sub>2</sub><sup>-</sup>, N<sub>2</sub>O |
| G | NO<sub>3</sub><sup>-</sup>, NO<sub>2</sub><sup>-</sup>, N<sub>2</sub>O | NO<sub>3</sub><sup>-</sup>, NO<sub>2</sub><sup>-</sup>, N<sub>2</sub>O |

The MLVSS concentration was measured in triplicate before the start of each batch test and all batch tests were carried out in duplicate. Apparent biomass-specific reduction rates for all nitrogen oxides, electron consumption and distribution rates were calculated in duplicate as outlined in previous studies (SI, Appendix 2). Detailed analytical methods for measuring phosphate, ammonium, nitrite, nitrate, VFA, PHA and glycogen have been outlined in the Supplementary Information (Appendix 1). GraphPad Prism (v6) was used for all statistical analyses.

### 2. 3. DNA extraction, sequencing library preparation, 16S amplicon sequencing and analysis

For microbial community analysis, 2 mL of mixed liquor from the three lab-scale reactors was collected on day 0 and day 28, coinciding with the first and last day of the batch tests. The genomic DNA extraction protocol can be found in the SI, Appendix 1. The library pool was sequenced at the Singapore Centre for Environmental Life Sciences Engineering (SCELSE) sequencing facility on a MiSeq (Illumina, US) using MiSeq Reagent kit V3 (2×300 paired end). Sequenced sample libraries were processed according to published DADA2 pipelines using the *dada2* R package (v 1.14). Details of the raw reads processing and community analysis are also explained in details in the SI, Appendix 1.

### 2.4 Model details

It is well established that during the anoxic phase in DPR systems, nitrate, nitrite or nitrous oxide are utilized as terminal electron acceptor by Accumulibacter for replenishment of its polyphosphate base, growth and other metabolic activities. The utilization rate of these terminal electron acceptors in Accumulibacter is associated with the PHA consumption rate for anoxic growth and subsequent phosphorus uptake. Thus, each step of the denitrification pathway (from NO_3_^−^ to N_2_ via NO_2_^−^, NO and N_2_O) can be associated with anoxic growth and phosphorus uptake rates. The model developed by Liu et al. (2015) incorporates these features to evaluate electron competition and nitrous oxide accumulation during denitrifying phosphorus removal [35]. In this study, we extended the application of the model by considering the simultaneous availability of multiple electron acceptors.

Each sequentially occurring denitrification step was associated with individual reaction-specific kinetics, i.e., anoxic growth rate on nitrate (μ_DPAO1_), nitrite (μ_DPAO2_), and nitrous oxide (μ_DPAO4_) and their associated phosphorus uptake rates. Experimental data from the batch tests were used to calibrate the relevant reactions and relationships for Accumulibacter enrichments, polyphosphate (X_PP_), PHAs (X_PHA_), DPAOs (X_DPAO_), residual inert biomass (X_I_) and seven soluble compounds - phosphate (S_PO4_), nitrate (S_NO3_), nitrite (S_NO2_), nitric oxide (S_NO_), nitrous oxide (S_N2O_), nitrogen gas (S_N2_) and readily degradable substrate (S_s_). The saturation constants for polyphosphate storage (K_PP,DPAO_, K_max,DPAO_) and PHA storage (K_PHA_) incorporated the relative abundance of DPAOs as assessed by 16S rRNA amplicon sequencing. The stoichiometry, component definition, model fitting parameters for Accumulibacter_nitrite_ and Accumulibacter_nitrate_, composition matrix and kinetic rate expression matrix and literature references are presented in the SI, Appendix 3.

## 3. RESULTS AND DISCUSSION

### 3.1 Reactor performance and microbial community characterization

Nutrient concentration measurement and microbial community analysis of the enrichments were conducted regularly. 16S rRNA gene amplicon sequencing was followed by classification of the most abundant ASVs at the highest level of resolution using the SILVA database to determine the degree of enrichment of target organisms and assess whether the community had undergone major shifts during the batch tests (Figure 1). The community structure was complex in all enrichments, although a significant proportion of the microbial community consisted of target organisms along with a low abundance of known competitor organisms. Accumulibacter_nitrate_ and Accumulibacter_nitrite_ had 48 ± 4% and 39 ± 3% relative abundance of target organisms, respectively, and a comparatively lower abundance of Competibacter (< 4% relative abundance). The Competibacter enrichment during the batch tests had a relative abundance of 44 ± 4% target organisms and 6 ± 3% of Accumulibacter. Organisms not known to display PAO or GAO phenotypes detected in the reactors were related to *Thiothrix, Thiobacillus, Hyphomicrobium, SJA-28, Denitratisoma, Cyanobacteria* and *Terrimonas*; they were approximately fourfold less abundant than target organisms.

**Figure 1.**
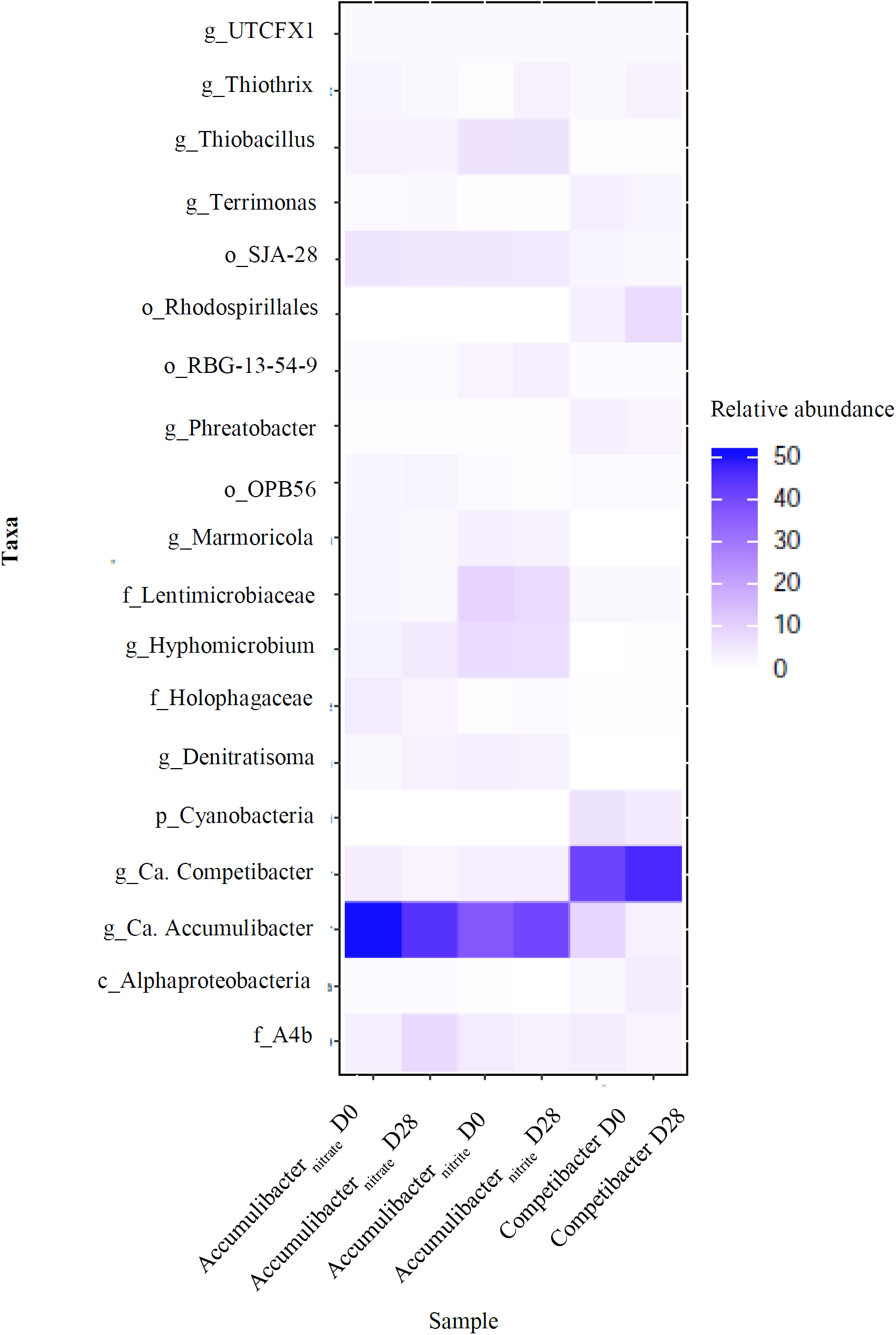
Microbial community analysis for samples collected at the start (D0) and end (D28) of batch tests from Accumulibacter_nitrite_, Accumulibacter_nitrate_, and Competibacter enrichment. Shown are the 15 most abundant organisms based on 16S rRNA gene amplicon sequencing and classified by the highest level of resolution obtained for each V1-V3 ASV annotated using the SILVA database. The microbial community was dominated by the respective target organism for each reactor and the abundance remained stable during the course of the experiment.

After an acclimation period lasting one month, pseudo-steady state activity and nitrogen removal was observed for Accumulibacter_nitrite_, Accumulibacter_nitrate_ and Competibacter-enriched sludge. During the anaerobic feeding phase, VFA was completely consumed followed by an increase in intracellular PHA content in all enrichments (Figure 2). Nitrogen oxides were utilized as terminal electron acceptors with concomitant phosphorus uptake in Accumulibacter_nitrite_ and Accumulibacter_nitrate_ enrichments. In the anoxic phase, Accumulibacter_nitrite_ showed 99% nitrite removal efficiency along with an anoxic phosphorus removal efficiency of 20%, whereas Accumulibacter_nitrate_ showed 72% nitrate removal efficiency with an anoxic phosphorus removal efficiency of 10%. The phosphorus remaining at the end of the anoxic phase in both enrichment reactors was utilized rapidly in the subsequent aerobic phase (Figure 2a, b). For Competibacter, the reducing power for PHA formation was obtained through glycolysis; therefore, glycogen reduction was observed with a concomitant increase in intracellular PHA in the anaerobic phase (Figure 2c). In the subsequent anoxic phase, nitrate added to the reactors was utilized to replenish the intracellular glycogen levels. The phosphorus concentration did not change during the entire cycle and remained below 1.5 mg/L.

**Figure 2.**
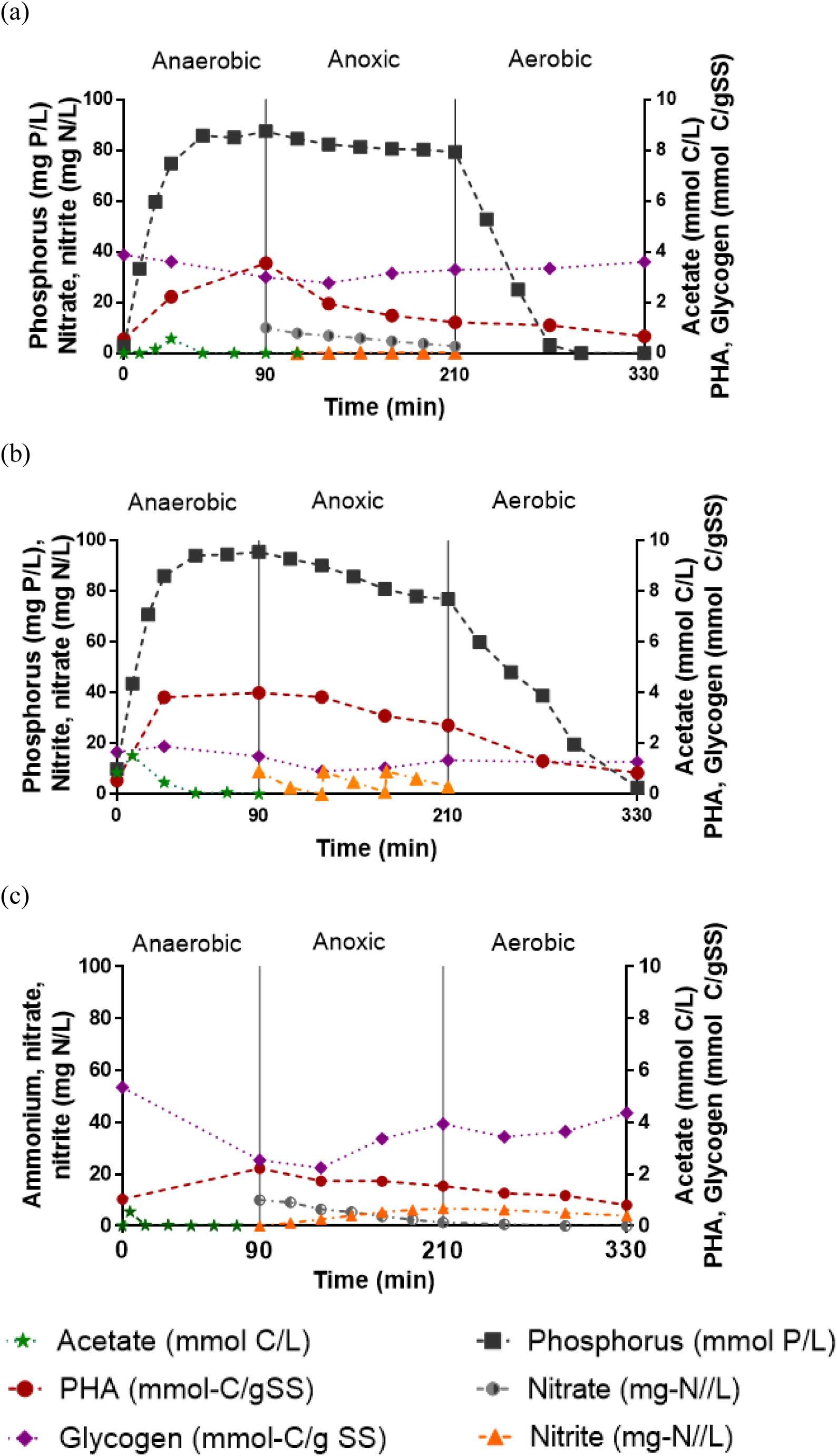
Transformation of nutrients and intracellular storage compounds during a typical SBR cycle of (a) Accumulibacter_nitrate_, (b) Accumulibacter_nitrite,_ and (c) Competibacter.

The microbial community of Accumulibacter enrichments utilized in previous studies included a significant fraction of GAOs, which could have led to confounding factors in understanding the denitrification kinetics of the target organism [26, 35-38]. In this study, we performed batch studies with Accumulibacter and Competibacter enrichments dominated by the target organisms to improve our understanding of their true denitrifying abilities.

### 3.2 Denitrification kinetics with single terminal electron acceptors

To test the denitrification kinetics and propensity for N_2_O accumulation in Accumulibacter_nitrate_, Accumulibacter_nitrite_ and Competibacter enrichments, true nitrous oxide, nitrate and nitrite reduction rates were measured in batch tests A, B and C, respectively (Figure 3a, b, c). These rates were calculated from the observed nitrogen oxide (NO_x_) concentration during each test (SI, Appendix 2). A linear decrease in NO_x_ concentrations was observed in tests with a single terminal electron acceptor, indicating denitrification activity (Fig S1).

**Figure 3.**
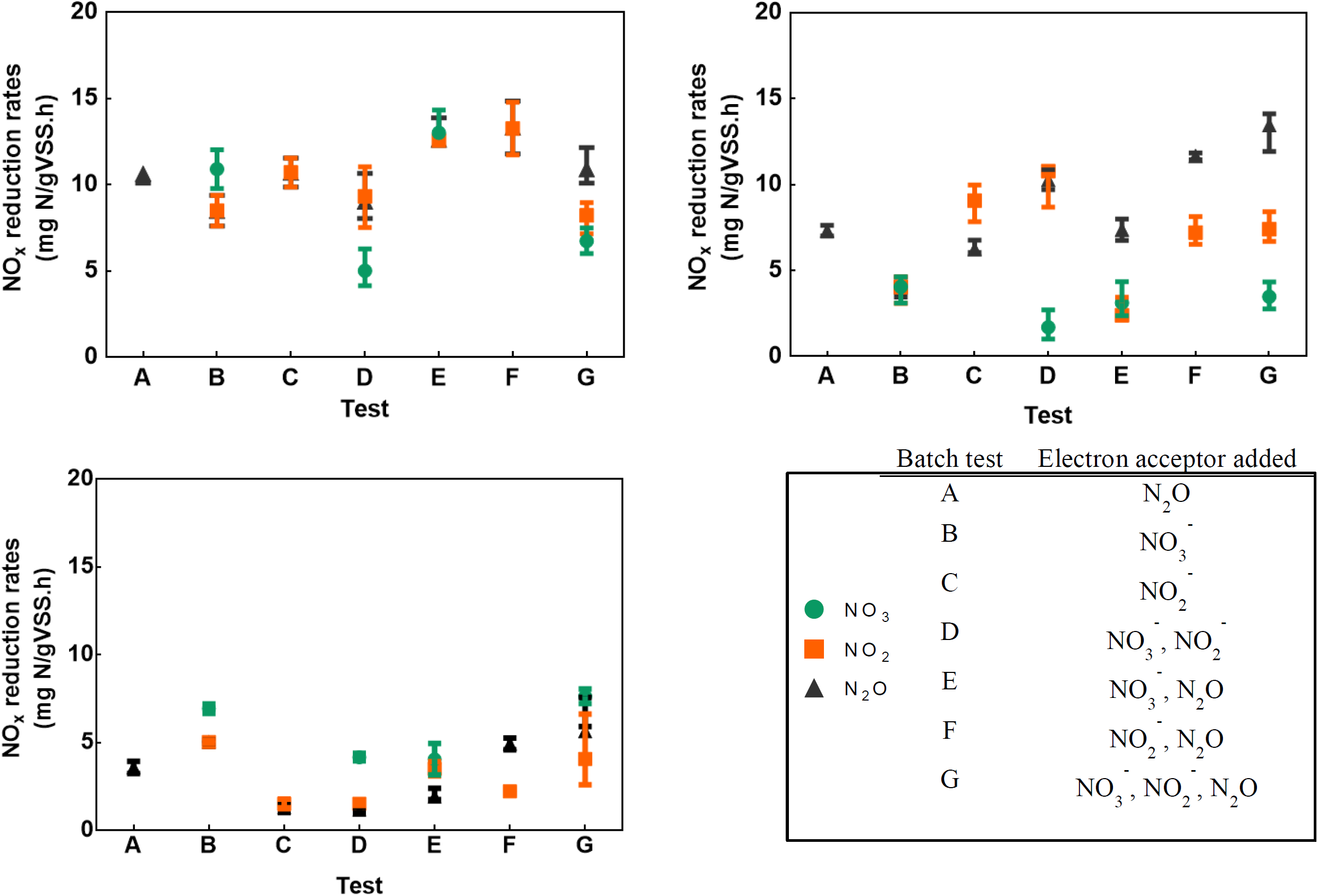
True reduction rate of nitrate, nitrite and N_2_O in batch test conditions A to G for (a) Accumulibacter_nitrate_ (b) Accumulibacter_nitrite_, and (c) Competibacter enrichments. The mean and range shown here were obtained from two replicates.

In batch test A, nitrous oxide reduction rates for Accumulibacter_nitrite_ and Accumulibacter_nitrate_ enrichments were 7.3 ± 0.43 and 10.6 ± 0.007 mg N gVSS^-1^ h^-1^, respectively. These rates were significantly higher than that of Competibacter (p < 0.05). Previously, it had been hypothesized that denitrification using PHA leads to N_2_O emissions [25, 39]. Yet, neither experimental nor modelling studies found any evidence to support this claim, suggesting other factors such as pH, free nitrous acid concentration and excess aeration as causes for increased N_2_O emissions [26, 34, 40, 41]. It is interesting that we observed higher nitrous oxide (N_2_O) utilization rates by Accumulibacter than by Competibacter, despite similar PHA storage and anoxic PHA utilization rates (Table 4). This rules out the possibility of incomplete denitrification and nitrous oxide accumulation solely due to PHA serving as carbon source [13].

For batch test B, nitrate reduction rates for Accumulibacter_nitrate_ and Competibacter were 10.9 ± 0.3 and 5.03 ± 0.39 mg N gVSS^-1^ h^-1^, respectively. These reduction rates were significantly higher (p < 0.05) than that of Accumulibacter_nitrite_, indicating that Accumulibacter acclimated to nitrite was less efficient in reducing nitrate. Also, as nitrate was reduced, nitrite accumulation was observed only for the Competibacter enrichment. In batch test C, the nitrite reduction rates for Accumulibacter_nitrite_ and Accumulibacter_nitrate_ were 9.1 ± 0.8 and 10.7 ± 1.2 mg N gVSS^-1^ h^-1^, respectively, which was almost six times higher than that observed for Competibacter. The preferential utilization of nitrate by Competibacter with concomitant nitrite accumulation was also observed in cycle studies, where the anoxic reaction phase was longer than in the batch test (Figure 1c). These differences in nitrite reduction between the enrichments support the conclusion that Competibacter had a lower preference for nitrite than Accumulibacter (p < 0.05).

Previously, McIlroy et al. (2015) reported genomes of two Competibacter GAOs (*Ca.* Competibacter denitrificans and *Ca.* Contendobacter odensis) encoding different denitrification pathways [7]. *Ca.* denitrificans encodes the complete denitrification pathway, while *Ca.* odensis only encodes genes for nitrate to nitrite reduction. Another study provided evidence of *nos* gene expression in denitrifying communities dominated by *Ca.* Competibacter denitrificans, signifying its active role in N_2_O reduction [27]. Consistent with our physiological data, the Competibacter enrichment was dominated by members of a Competibacter-lineage that could predominantly reduce nitrate to nitrite, and showed limited N_2_O uptake despite adequate PHA storage. Genomic analyses of various Accumulibacter clades have revealed that most members encode the *nos* gene, and have the potential to utilize N_2_O as an electron acceptor [16]. This was reflected in batch tests where adequate N_2_O reduction was observed in Accumulibacter enrichments. Thus, in addition to adequate environmental conditions required to drive N_2_O reduction, enzymatic regulation of the *nos* gene is important in PHA-driven denitrification.

### 3.3 Denitrification kinetics with multiple terminal electron acceptors

Multiple electron acceptors were simultaneously added in batch tests D to G, and the observed NO_x_ reduction rates were compared with those from the single electron acceptor batch experiments (tests A, B and C). The tests allowed us to investigate the potential for electron competition limiting N_2_O utilization with different combinations of terminal electron acceptors for Accumulibacter and Competibacter. The overall NO_x_ reduction rates for the Accumulibacter and Competibacter enrichments increased in the presence of multiple electron acceptors, and were not affected by the simultaneous addition of nitrogen oxides (Figure 3). In a previous study, another phosphorus accumulating organism, *Tetrasphaera*, had shown reduced denitrification kinetics when multiple electron acceptors were added simultaneously, indicating a limited electron supply to different denitrification enzymes [42]. In this study, we observed the opposite with an increase in overall NO_x_ reduction rates in the presence of multiple electron acceptors, suggesting the absence of electron competition.

In batch test D, when nitrite and nitrate were added together as terminal electron acceptors, the nitrate reduction rate was significantly higher for Accumulibacter_nitrate_ compared to Accumulibacter_nitrite_ (p < 0.0005). For both Accumulibacter enrichments, the net nitrite reduction rate was observed to be two to five times higher than the net nitrate reduction rate, with no observed N_2_O accumulation, suggesting that the Accumulibacter taxa enriched in both reactors were able to take up nitrite provided externally as an electron acceptor, despite the simultaneous presence of nitrate. By comparison, for Competibacter, the true nitrate reduction rate was twice the true nitrite reduction rate, leading to accumulation of nitrite (Figure 3). The Competibacter enrichment displayed preferential utilization of nitrate despite the simultaneous presence of equimolar amounts of nitrate and nitrite, reinforcing our observations from the batch tests of its limited nitrite utilization ability.

In batch test E, when nitrate and N_2_O were utilized as terminal electron acceptors, the N_2_O utilization rate for Accumulibacter_nitrite_ was twice its nitrate utilization rate. In contrast, for Accumulibacter_nitrate_, the nitrate, nitrite and N_2_O reduction rates were similar, and complete denitrification was observed. However, unlike Accumulibacter_nitrite_ the externally provided N_2_O was not utilized despite similar PHA levels. This is also reflected in the electron consumption profiles for batch test E (Figure 4), where Accumulibacter_nitrate_ showed an equal fraction of electrons consumed during each denitrification step, while in Accumulibacter_nitrite_ most electrons were consumed for nitrate and N_2_O reduction. When nitrite and N_2_O were added together (batch test F, Figure 4), Accumulibacter_nitrite_ showed N_2_O uptake in excess of that expected by denitrification of nitrite; by comparison, Accumulibacter_nitrate_ exhibited similar nitrite and N_2_O reduction rates with no excess N_2_O uptake. Finally, in batch test G where nitrate, nitrite and N_2_O were added together, the total denitrification rate for both Accumulibacter enrichments was 1.5 to 2 times higher than that of Competibacter.

**Figure 4.**
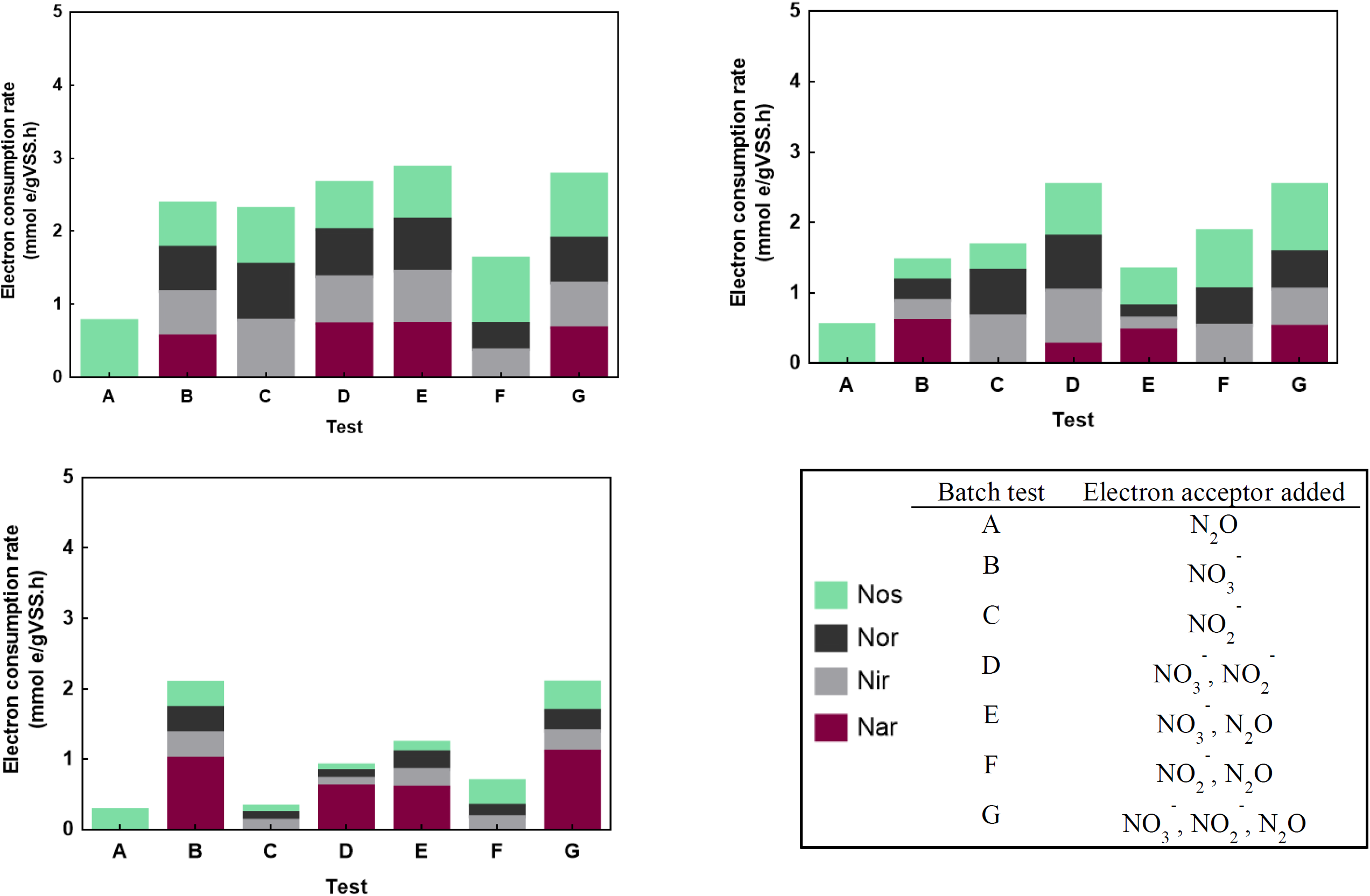
Electron consumption rates by specific denitrification enzymes for batch test conditions A to G for (a) Accumulibacter_nitrate_, (b) Accumulibacter_nitrite_, and (c) Competibacter. Denitrification enzymes are nitrate reductase (Nar, shown in maroon), nitrite reductase (Nir, in grey), nitric oxide reductase (Nor, in black) and nitrous oxide reductase (Nos, in green).

In comparison to previous studies measuring denitrification kinetics in DPAOs and DGAOs, nitrous oxide accumulation was not observed in this study when multiple electron acceptors were added simultaneously (Table 2) [36, 43]. FNA concentrations in previous studies ranged from 0.001 to 0.015 mg HNO_2_-N L^-1^, which is much higher than the suggested inhibitory concentrations of 0.0007 – 0.001 mg HNO_2_-N L^-1^ [26]. FNA in excess of inhibitory concentrations has been shown to affect the *nos* gene transcriptional process, and directly react with the copper-containing active sites of N_2_O reductase (Nos) [44, 45]. Therefore, in this study, the pH and NOx dosage study were adjusted such that the FNA formed was below the inhibitory levels (0.1 – 0.2 µg HNO_2_-N L^-1^). It has also been proposed that the limited bioenergetic advantage of N_2_O reduction for a cell reducing nitrate to nitrogen gas (i.e., ∼20% of the total energy generated) allows for release of N_2_O as the final denitrification product instead of N_2_ [29]. Further, utilizing N_2_O produced within a cell generates more energy than transporting N_2_O into the cell and subsequently reducing it to nitrogen gas [42]. However, in this study, we observed that Accumulibacter was able to utilize N_2_O that was externally provided in addition to N_2_O produced as a result of denitrification of NO_x_. The electron distribution patterns for Accumulibacter, too, indicated a significant portion of electrons was consumed by Nos, which was not limited by the simultaneous presence of other NO_x_ (SI, Fig S3). Although the gene expression levels of various denitrification enzymes involved were not measured in this study, previous studies have identified a strong correlation between the transcription of *nosZ* genes (clade I and II) and the N_2_O reduction rate [10, 27].

**Table 2.** Comparison of the amount of N_2_O accumulated per N reduced (in mg/L) with values in previous studies

| Study | FNA level<br>calculated<br>(HNO<sub>2</sub>-N. L<sup>-1</sup>) | Batch test (electron acceptor added) |  |  |  |  |  |
| --- | --- | --- | --- | --- | --- | --- | --- |
|  |  | Test B<br>(NO<sub>3</sub><sup>-</sup>) | Test C<br>(NO<sub>2</sub><sup>-</sup>) | Test D<br>(NO<sub>2</sub><sup>-</sup> & NO<sub>3</sub><sup>-</sup>) | Test E<br>(NO<sub>3</sub><sup>-</sup> & N<sub>2</sub>O) | Test F<br>(NO<sub>2</sub><sup>-</sup> & N<sub>2</sub>O) | Test G<br>(NO<sub>2</sub><sup>-</sup>, NO<sub>3</sub><sup>-</sup> & N<sub>2</sub>O) |
| **Acc<sup>a</sup>** | 0.1 - 0.2 µg | 0.31 ± 0.01 | 6.93 ± 0.38 | 3.63 ± 0.18 | - | - | - |
| **Acc<sup>b</sup>** | 0.1 - 0.2 µg | - | 30.97 ± 1.06 | 8.40 ± 0.01 | - | - | - |
| **Comp<sup>c</sup>** | 0.1 - 0.2 µg | - | 15.83 ± 2.59 | 1.07 ± 0.97 | 1.57 ± 0.26 | - | - |
| **Tet<sup>d</sup>** | 5 µg | 16.7 ± 0.8 | 15.5 ± 0.8 | 17.7 ± 0.9 | - | - | - |
| **DPAO<sup>e</sup>** | 1.58 µg | 8.72 ± 0.2 | 17.4 ± 5.9 | 31.2 ± 2.70 | - | 20.11 ± 1.90 | 11.30 ± 3.10 |
| **DGAO<sup>f</sup>** | 1.58 µg | 7.12 ± 2.16 | 82.95 ± 4.79 | 45.45 ± 0.89 | 13.71 ± 5.81 | 56.90 ± 4.92 | 48.45 ± 5.94 |

<sup>a</sup> *Ca. Accumulibacter* (enriched with nitrate as terminal electron acceptor in this study) (this study)
<sup>b</sup> *Ca. Accumulibacter* (enriched with nitrate as terminal electron acceptor in this study) (this study)
<sup>c</sup> *Ca. Competibacter* (enriched with nitrate as terminal electron acceptor in this study)
<sup>d</sup> *Tetrasphaera* culture (Marques et al. 2018)
<sup>e</sup> DPAO culture (*Accumulibacter* clade I and II present) (Ribera et al., 2016)
<sup>f</sup> DGAO culture (*Competibacter* and *Defluviicoccus* present) (Ribera et al., 2016)
- Below detection limit

**Table 3.** Mean PHA concentration and standard deviation at the beginning and end of batch tests

| Enrichment | Phase | Batch test (electron acceptor added) |  |  |  |  |  |  |
| --- | --- | --- | --- | --- | --- | --- | --- | --- |
|  |  | Test A<br>(N<sub>2</sub>O) | Test B<br>(NO<sub>3</sub><sup>-</sup>) | Test C<br>(NO<sub>2</sub><sup>-</sup>) | Test D<br>(NO<sub>2</sub><sup>-</sup> & NO<sub>3</sub><sup>-</sup>) | Test E<br>(NO<sub>3</sub><sup>-</sup> & N<sub>2</sub>O) | Test F<br>(NO<sub>2</sub><sup>-</sup> & N<sub>2</sub>O) | Test G<br>(NO<sub>2</sub><sup>-</sup>, NO<sub>3</sub><sup>-</sup> & N<sub>2</sub>O) |
| Accumulibacter<sub>nitrate</sub> | Start (90 min) | 10.59 ± 1.58 | 7.16 ± 1.09 | 7.37 ± 0.16 | 8.60 ± 0.80 | 8.74 ± 1.02 | 9.14 ± 0.01 | 6.07 ± 0.39 |
|  | End (120 min) | 9.54 ± 0.57 | 5.71 ± 1.13 | 6.09 ± 0.08 | 6.13 ± 0.10 | 4.77 ± 0.82 | 6.64 ± 0.71 | 3.80 ± 0.23 |
| Accumulibacter<sub>nitrite</sub> | Start (90 min) | 6.57 ± 0.44 | 10.06 ± 0.33 | 10.75 ± 0.97 | 9.97 ± 0.07 | 10.15 ± 0.31 | 13.88 ± 0.34 | 7.64 ± 0.51 |
|  | End (120 min) | 5.67 ± 0.96 | 9.43 ± 0.88 | 9.18 ± 1.60 | 8.89 ± 0.15 | 9.07 ± 0.52 | 12.67 ± 0.60 | 7.21 ± 0.49 |
| Competibacter | Start (90 min) | 5.03 ± 0.91 | 8.16 ± 1.06 | 6.23 ± 0.34 | 9.97 ± 1.14 | 7.96 ± 0.29 | 6.16 ± 0.16 | 8.16 ± 0.07 |
|  | End (120 min) | 4.04 ± 1.21 | 7.66 ± 0.32 | 6.52 ± 0.08 | 8.12 ± 0.76 | 6.19 ± 0.15 | 5.63 ± 0.02 | 6.36 ± 0.14 |

It is likely that a combination of factors such as adequate intracellular PHA storage, higher availability of N_2_O and low FNA inhibition in this study allowed for adequate *nos* transcription. This made it bioenergetically feasible for Accumulibacter to transport and reduce externally added N_2_O, while simultaneously utilizing N_2_O produced by nitrate or nitrite reduction.

### 3.4 Electron consumption and distribution during denitrification

For the batch tests conducted, we compared electron consumption and distribution trends as they have been previously shown to be affected by electron competition [28]. Here, Accumulibacter enrichments were observed to have a higher electron consumption rate than Competibacter, which can be attributed to higher denitrification rates observed in the former. The total average electron consumptions in multiple electron addition tests for Accumulibacter_nitrite_ and Accumulibacter_nitrate_ were 2.1 ± 0.63 and 2.47 ± 0.58 mmol e^-^ g VSS^-1^ h^-1^, respectively (Figure 4). These values were higher than the total electron consumptions calculated for Accumulibacter_nitrite_ with nitrite addition (test C, 1.44 mmol e^-^ gVSS^-1^ h^-1^) and for Accumulibacter_nitrate_ with nitrate addition (test B, 2.36 mmol e^-^ gVSS^-1^ h^-1^). Thus, despite the simultaneous presence of multiple electron acceptors, the PHA-derived electron supply to denitrification enzymes increased as a function of the total NO_x_ concentration, demonstrating that intracellular PHA storage was sufficient to support denitrification in Accumulibacter.

Previously, it has been surmised that enzyme-specific affinity to electron carriers and regulation of electron transfer are important factors that determine electron distribution to enzymes involved in denitrification [28]. It is usually observed that denitrification enzymes upstream (such as Nar) have a greater ability to receive electron carriers than downstream enzymes (e.g. Nir and Nos) [46]. This is because the ubiquinone/ubiquinol pool supplies electrons to the Nar enzymes and the cytochrome c550/pseudoazurin pool, and the latter subsequently supplies electrons to the Nir and Nos enzymes. In this study, Accumulibacter_nitrate_ and Competibacter enrichments showed significantly higher electron consumption rates in batch test E (NO_3_^-^ and N_2_O added) compared to batch test F (NO_2_^-^ and N_2_O added) (p-value = 0.007 and 0.044 for Accumulibacter_nitrate_ and Competibacter, respectively). However, the trend for electron consumption was reversed for Accumulibacter_nitrite_ with electron consumption in batch test F approximately 1.5 times that in test E (Figure 4) (p-value = 0.028). Thus electron competition was not necessarily the limiting factor for denitrification, enrichment-specific affinity for different electron acceptors also affected the overall NO_x_ reduction. Further, in Accumulibacter_nitrite_ during nitrite reduction, 37% of total electrons were distributed to nitrite reductase whereas in the presence of other terminal electron acceptors, i.e., tests D, F and G, the fractions of total electrons distributed to nitrite reductase were reduced to 30%, 27%, and 22%, respectively. This proves that nitrite reductase did not display a higher ability to compete for electrons compared to nitrate or nitrous oxide reductase, as has been reported for heterotrophic organisms and some *Tetrasphaera*-related species [28, 42].

We also compared the phosphorus-uptake-to-electron-consumption ratio in all batch tests of Accumulibacter_nitrite_ and Accumulibacter_nitrate_, because the total electron consumption is indicative of the energy utilized for anoxic phosphorus uptake (Figure 5). This ratio, just as the true denitrification rate, did not seem to be affected by increasing concentrations of electron acceptors, but rather by the type of electron acceptor present. Phosphorus uptake associated with nitrate in Accumulibacter_nitrate_ enrichments was twice that observed for Accumulibacter_nitrite_. Higher phosphorus uptake was associated with nitrite and nitrous oxide compared to nitrate in Accumulibacter_nitrite_ enrichments, emphasizing the impact of long-term adaptation to certain terminal electron acceptors on observed denitrification kinetics and associated phosphorus uptake.

**Figure 5.**
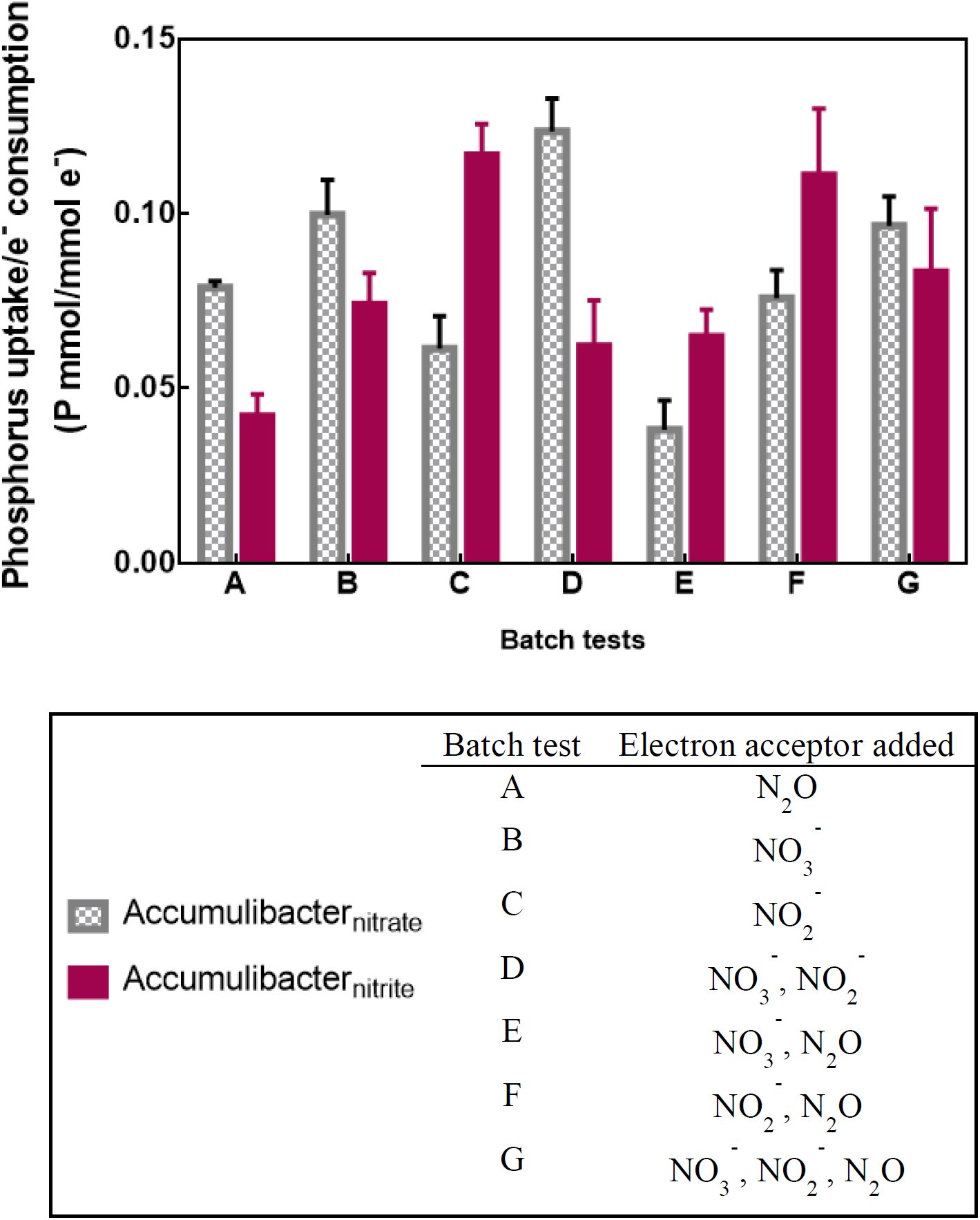
Ratio of P uptake per electron consumption (mmol e^-^) obtained for the batch tests performed with various terminal electron acceptors for the two Accumulibacter enrichment cultures – Accumulibacter_nitrite_ and Accumulibacter_nitrate_.

In summary, the analysis of electron consumption and distribution in this study leads to the conclusion that electron supply increased as a function of nitrogen oxide concentration, indicating that adequate PHA levels can drive denitrification in Accumulibacter. Upstream denitrification enzymes did not limit the flow of electrons to Nos, and hence there was no electron competition for nitrous oxide reduction in Accumulibacter. A comparison of the electron distribution in the two Accumulibacter enrichments suggests an enrichment-specific affinity for terminal electron acceptors.

### 3.5 Model evaluation of denitrification kinetics and electron competition

The nitrate, nitrite, and nitrous oxide reduction rates for Accumulibacter measured during the batch tests with low FNA inhibition were employed in a metabolic model to predict anoxic growth at each denitrification step and the associated phosphorus uptake for DPAOs [35]. The model used kinetic parameters specific for each step in the denitrification process, and calculated a carbon oxidation rate to predict the accumulation of denitrification products. As the flanking community members were much less abundant than the Accumulibacter cells present in the enrichments, their contribution to NO_x_ utilization was considered negligible. The model was optimized by recalibrating four key kinetic parameters: rate constant for storage of polyphosphate (q_PP_), anoxic growth rate on nitrate (μ_DPAO1_), anoxic growth rate on nitrite (μ_DPAO2_) and anoxic growth rate on N_2_O (μ_DPAO4_). In addition to these parameters, another 17 kinetic and 6 stoichiometric parameters were obtained from the literature (Table S1).

The optimized model captured the trends well for the reduction of various nitrogen oxides (NO_x_) and phosphorus uptake in the batch tests. The R^2^ values obtained for both Accumulibacter enrichments were 0.97 ± 0.01, 0.96 ± 0.09, 0.97 ± 0.05 and 0.96 ± 0.10 for nitrate, nitrite, N_2_O and phosphorus, respectively, confirming the robustness of the model and parameter values to accurately predict N_2_O accumulation in Accumulibacter enrichments (Figure 6). A comparison of the anoxic growth rates optimized with the model revealed that the growth on nitrite, μ_DPAO2_, was lower for Accumulibacter_nitrate_ compared to Accumulibacter_nitrite_, indicating differences in nitrite reducing abilities (p-value = 0.04). These observed differences also highlight the effect of long-term adaptation to certain terminal electron acceptors.

**Figure 6.**
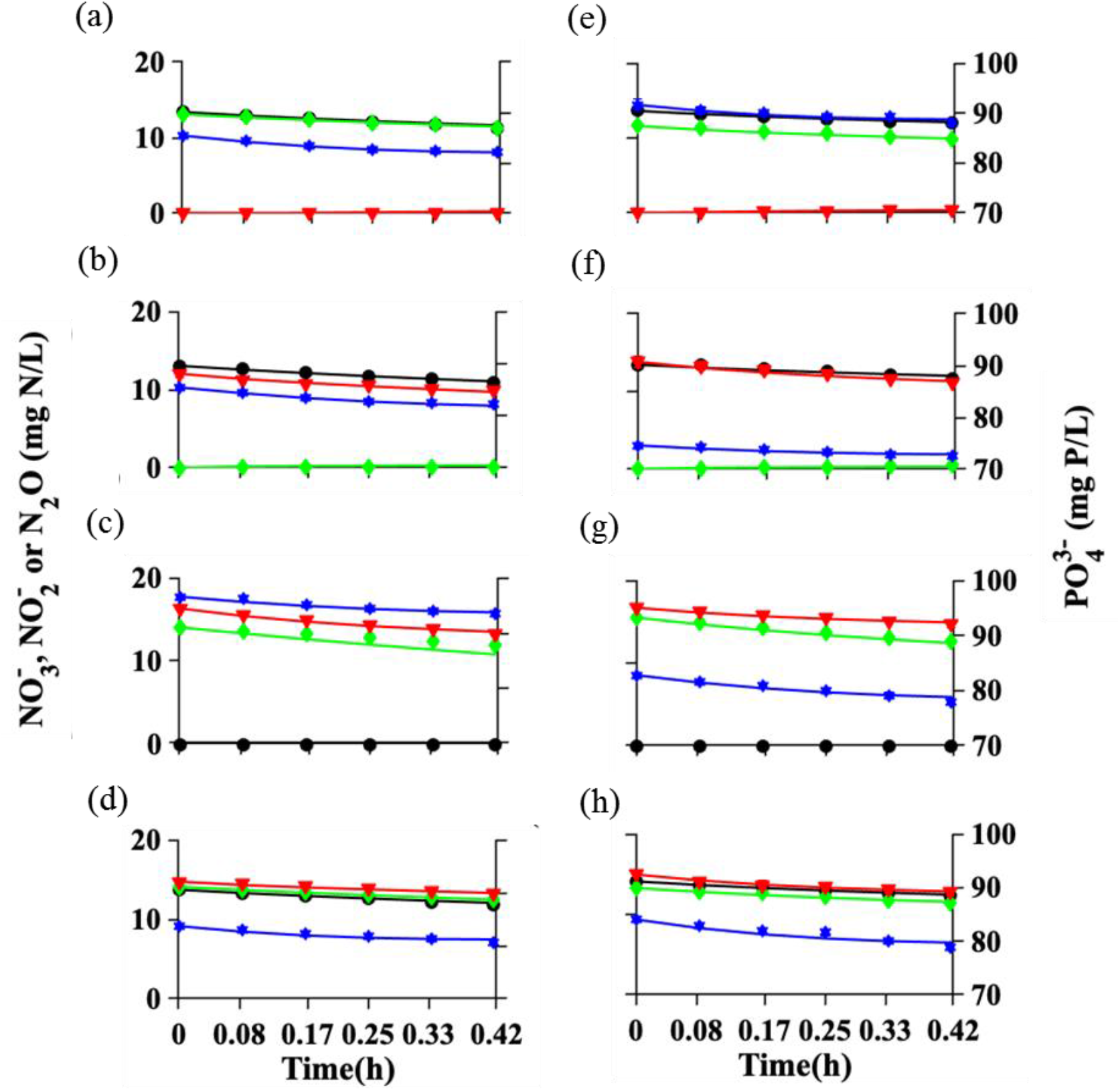
Simulation of batch perturbation studies of nitrate-enriched Accumulibacter with (a) NO_3_^−^ and NO_2_^−^, (b) NO_3_^−^ and N_2_O, (c) NO_3_^−^ and N_2_O, and (d)) NO_3_^−^, NO_2_^−^ and N_2_O. Simulation of batch perturbation study of nitrite-enriched Accumulibacter with (e) with) NO_3_^−^ and NO_2_^−^, (f)) NO_3_^−^ and N_2_O, (g) NO_2_^−^ and N_2_O, and (h) with) NO_3_^−^, NO_2_^−^ and N_2_O. x_axis limit (0 – 0.42), y_axis_left limit (0 – 20) and y_axis_right limit (70 – 90). Figure legend: predicted PO_4_^3-^ (**–**, solid blue line), predicted NO_3_^-^ (**–**, solid black line), predicted NO_2_^-^ (**–**, solid green line), predicted N_2_O (**–**, solid red line), measured PO_4_^3-^ (★), measured NO_3_^-^ (●), measured NO_2_^-^ (✦), and measured N_2_O (▼).

In batch tests where N_2_O was added in addition to nitrite or nitrate, Accumulibacter effectively utilized N_2_O that was produced from denitrification and, more importantly, did not lead to any N_2_O accumulation. A previous study reported a similar observation for *Tetrasphaera*-related organisms, suggesting that high availability of N_2_O and energy from nitrate/nitrite reduction in these tests led to an increased synthesis of nitrous oxide reductase (Nos) [42]. Here, we also used the model to evaluate energy supplied for Nos synthesis (SI, Appendix 3, Table S6). For Accumulibacter, the anoxic growth rate on N_2_O increased approximately 1.5 to 2 times when multiple electron acceptors were added simultaneously (tests D to G) as compared to tests with single terminal electron acceptors (tests A to C) (p-value = 0.03 and 0.0048 for Accumulibacter_nitrite_ and Accumulibacter_nitrate_, respectively). This further confirms our hypothesis that the increased availability of N_2_O and energy derived from the reduction of multiple electron acceptors allowed for the increase in anoxic growth rate.

Other mathematical models have been used to predict phosphorus removal, but few compared the nitrous oxide (N_2_O) accumulation from denitrifying phosphorus removal processes. A limitation is that the denitrification process is usually represented as a single-or two-step process. One study utilized a similar four-step denitrification model as our approach to predict nitrous oxide (N_2_O) accumulation during biological nitrogen removal in anaerobic/anoxic/oxic sequencing batch reactors (A^2^O-SBR), but abstained from accounting for the abundant Accumulibacter and Competibacter organisms usually found in these systems [47]. It can be surmised that lack of organism-specific parameters in the model can lead to a lower precision of N_2_O accumulation prediction. In contrast, the model applied in this study lacks the aforementioned shortcomings and adequately describes N_2_O accumulation during denitrification [35].

The model heuristic can also be integrated with other N_2_O models for nitrification and denitrification to gain more insight into N_2_O dynamics during various stages of biological wastewater treatment. FNA concentrations kept below inhibitory levels in this study allowed for modelling of the true denitrification kinetics in Accumulibacter. FNA inhibition leading to N_2_O accumulation is not limited to polyphosphate accumulating organisms, but has also been documented for heterotrophic, hydrogenotrophic, ammonia-oxidising and Anammox organisms [48-51]. A lower tendency for N_2_O accumulation and higher anoxic growth rate on N_2_O was observed when FNA concentrations were low. This identifies FNA as an important parameter that must be included in models developed to predict N_2_O accumulation based on nitrite concentration and pH. To supplement these modelling predictions, future studies can also include metatranscriptomic analysis of the denitrification enzymes in Accumulibacter, and compare levels of transcription at different FNA concentrations.

## 4. CONCLUSIONS

Denitrifying phosphorus removal (DPR) is a promising technology. In this work we have focussed on the potential for nitrous oxide accumulation by dominant microbial community members - Accumulibacter and Competibacter - in DPR systems. Our batch test results and validation with a metabolic model suggest that denitrification by Accumulibacter and Competibacter is not limited by electron competition. In fact in Accumulibacter, anoxic growth on nitrous oxide (N_2_O) in the presence of other nitrogen oxides was significantly higher compared to when nitrous oxide added as sole electron acceptor. Our observations indicate that sufficiently high PHA levels and low free nitrous acid (FNA) inhibition resulted in increased transcription of the *nos* gene, allowing for higher N_2_O reduction than previously observed. The denitrification kinetics for Competibacter revealed poor nitrite or nitrous oxide utilization despite sufficient PHA storage and low FNA inhibition, implying a truncated denitrification pathway. To conclude, the reduction of nitrogen oxides by Accumulibacter and Competibacter is largely governed by their denitrification enzyme-specific affinity to electron carriers, the availability of sufficient internal storage polymers to provide energy for electron transfer and low concentrations of FNA. Understanding the nitrous oxide reduction potential of the dominant microbial community members in DPR can be leveraged to design energy efficient treatment processes for biological phosphorus removal and subsequently reduce unwanted emissions.

## Supporting information

Roy et al. 2020

## AUTHOR CONTRIBUTIONS

The study was designed by SR, SW and HYN. SR conducted the batch test experiments and analysed the microbial community. SR, NP and SW analysed and interpreted the data obtained over the course of this study and wrote the manuscript with contributions from HYN.

## ACKNOWLEDGEMENTS

This research was supported by the Ministry of Education, Singapore under the Research Centre of Excellence Programme. We acknowledge Dr. Daniela Drautz-Moses for performing 16S rRNA gene amplicon sequencing. Many thanks go to Dr. Rohan Williams for his inputs on interpretation of the sequencing data and for proofreading the manuscript. We also extend our gratitude to James Jun, Muhamad Danial Bin Suthree, Ahmadul Hafiz Ain Azman Jun, Eganathan Kaliyamoorthy and Sara Swa Thi for performing nutrient tests during the course of this study.

