## Supplementary material for "Denitrification kinetics indicates nitrous oxide uptake is unaffected by electron competition in Accumulibacter": Roy et al. 2020

[a]Corresponding author:

**Appendix 1**

1. Analytical methods
2. DNA extraction, sequencing library preparation, 16S amplicon sequencing and analysis

**Appendix 2**

1. $N_2O$ measurement and calculation

**Appendix 3**

1. Observed $NO_x$ reduction rates
2. Electron distribution profile
3. Kinetic and stoichiometric parameters for DPAOs cultures grown with $NO_3^-$ and $NO_2^-$.
4. Kinetic rate expressions for the DPAO model
5. Peterson matrix for the DPAO model
6. Model fitting parameters for $Accumulibacter_{nitrate}$ and $Accumulibacter_{nitrite}$
7. Best fit parameters for anoxic growth rate

**Appendix 1**

1. *Analytical methods*

To monitor reactor performance and nutrient profile, mixed liquor samples were collected regularly from reactors for phosphate, ammonium, nitrite, nitrate and volatile fatty acids (VFA) analyses. Phosphate, ammonium, nitrite and nitrate were analyzed using ion chromatography (Prominence, Shimadzu) fitted with Shim-pack IC-SA2 (250 mm length, 4.0 mm ID) for anions and Shim-pack IC-C4 (150 mm length, 4.6 mm ID) for cations. VFA concentrations (acetate, propionate, butyrate) were analyzed using gas chromatography (Prominence, Shimadzu) fitted with a flame ionization detector (FID) fitted with a DB-FFAP (30 m length, 0.25 m diameter, and 0.25 µm film) column (Agilent Technology, USA). Glycogen and PHA extraction protocols were optimized for granular sludge as per Lanham et al [1, 2]. Glycogen was measured using high performance liquid chromatography (Prominence, Shimadzu) equipped with a UV-Vis detector. PHA analysis was conducted using gas chromatography (Prominence, Shimadzu) equipped with a flame ionization detector (FID) and fitted with a DB-5MS Ultra Inert (30 m length, 0.25 m diameter, and 0.25 µm film) column (Agilent Technology, USA).

2. *DNA extraction, sequencing library preparation, 16S amplicon sequencing and analysis*

Genomic DNA was extracted from the mixed liquor samples using FastDNA™ 2mL SPIN Kit for Soil (MP Biomedicals, CA, USA) following an optimized protocol outline [Albertsen et al 2016]. Bacterial 16S rRNA gene amplicon sequencing was performed using V1-V3 region primer set (27F AGAGTTTGATCCTGGCTCAG and 534R ATTACCGCGGCTGCTGG). The 25 µL PCR matrix contained 2 mU Platinum

R Taq DNA polymerase high fidelity, 400 nm dNTPs, 1.5 mM $MgSO_4$, 1 Platinum R High Fidelity buffer (Thermo Fischer Scientific, USA), 10 ng genomic DNA, and a pair of barcoded library adaptors (400 nM). The thermoycler settings included 95°C for 2 min, 30 cycles of 95°C for 20 s, 56°C for 30 s, 72°C for 60 s and final elongation at 72°C for 5 min. PCR reactions were run in duplicates and pooled afterwards. The amplicon libraries were purified using the Agencourt R AMpure XP bead protocol (Beckmann Coulter, USA) with 1.8-bead solution/PCR solution ratio and the quality of DNA before submitting for sequencing was checked on the Agilent 2200 TapeStation. Based on amplicon sizes and library concentrations that were checked using Thermo Scientific Nanodrop™ 2000 and Invitrogen Qubit® 2.0, the samples were pooled in equimolar concentrations.

Sequenced sample libraries were processed according to published DADA2 pipelines using the *dada2* R package. This method was chosen because exact sequence analysis has been shown to have several benefits over traditional OTU clustering methods. Sequences were truncated at Phred scores of 30 at 290 and 260 nucleotides for forward and reverse reads, respectively. Truncation of reads with error rates higher than 5 and 6 in forward and reverse reads, respectively, were removed. From these reads, error rates was estimated, followed by sample inference, denoising, and finally merging forward and reverse reads by at least 12 bases. The resultant sequence table was then subjected to chimera removal and taxonomy assignment using the SILVA database (v.132). An abundance table generated from this analysis reflected the total count of each ASV detected in each sample. The counts in the abundance table were transformed to relative abundance with respect to the total reads in the samples. The phyloseq and ggplot R-packages were used for microbial community analysis and heatmap generation.

**Appendix 2**

1. *$N_2O$ measurement and calculation*

Dissolved $N_2O$ concentrations were monitored with an online $N_2O$ microsensor ($N_2O$ -R, Unisense A/S, Denmark). Total $N_2O$ generation ($r_g$) was determined according to equation 4-1, where $r_e$ was the $N_2O$ emission rate from water to air (mg N/L·s) measured in control tests and $r_a$ was the dissolved $N_2O$ accumulation rate obtained from the microsensor immersed in the mixed liquor (mg N/L·s). For the $N_2O$ emission rate from water to air ($r_e$), four control tests were carried out in duplicate and the averaged value of $N_2O$ loss due to stripping was used in equation (1) to estimate the total $N_2O$ generated during the course of the experiment.

$$\boldsymbol{r_g = r_e + r_a} \qquad \text{Equation 1}$$

Apparent biomass-specific reduction rates for all nitrogen oxides were determined by linear regression profiles of nitrate, nitrite and N2O profiles divided by VSS concentration ($r_{NO_3^-,a}$ , $r_{NO_2^-,a}$ , $r_{N_2O,a}$, respectively). For calculations, any accumulation of these compounds was represented as negative consumption. True reduction rate for each denitrification step was calculated as shown below where, $r_{NO_3^-}$ , $r_{NO_2^-}$ , $r_{NO}$and $r_{N_2O}$ are true reduction rates for nitrate, nitrite, nitric oxide and nitrous oxide, respectively (expressed as mg N/gVSS·h).

$$\boldsymbol{r_{NO_3^-} = r_{NO_3^-,a}} \qquad \text{Equation 2}$$

$$\boldsymbol{r_{NO_2^-} = r_{NO_2^-,a} + r_{NO_3^-}} \qquad \text{Equation 3}$$

$$\boldsymbol{r_{NO} = r_{NO,a} + r_{NO_2^-}} \qquad \text{Equation 4}$$

$$\boldsymbol{r_{N_2O} = r_{N_2O,a} + r_{NO}} \qquad \text{Equation 5}$$

As nitric oxide (NO) is known to be a potential cytotoxin, intracellular concentrations of nitric oxide are usually maintained at very low levels, which are possible only when the rate of nitric oxide formation equals the rate of nitrous oxide formation.

Thus NO reduction is prioritized by cells and hence is not a rate-limiting step in denitrification. Electron consumption rate for each denitrification step was calculated as follows:

$$\boldsymbol{r_{Nar}} = \frac{r_{NO_3^-}}{14} \times \mathbf{2} \qquad \text{Equation 6}$$

$$\boldsymbol{r_{Nir}} = \frac{r_{NO_2^-}}{14} \times \mathbf{1} \qquad \text{Equation 7}$$

$$\boldsymbol{r_{Nor}} = \frac{r_{NO}}{14} \times \mathbf{1} \qquad \text{Equation 8}$$

$$\boldsymbol{r_{Nos}} = \frac{r_{N_2O}}{14} \times \mathbf{1} \qquad \text{Equation 9}$$

where, $r_{Nar}$, $r_{Nir}$, $r_{Nor}$ $and$ $r_{Nos}$ are electron consumption rates for Nar, Nir, Nor and Nos (mmol e/g VSS·h) [3]. Electron distribution towards each step of $NO_x$ reduction was calculated as outlined by Ribera-Guardia et al. [4]:

$$\text{Electron distribution (\%)} = \frac{r_{NOx}}{r_{Nar} + r_{Nir} + r_{Nor} + r_{Nos}} \times 100 \qquad \text{Equation 10}$$

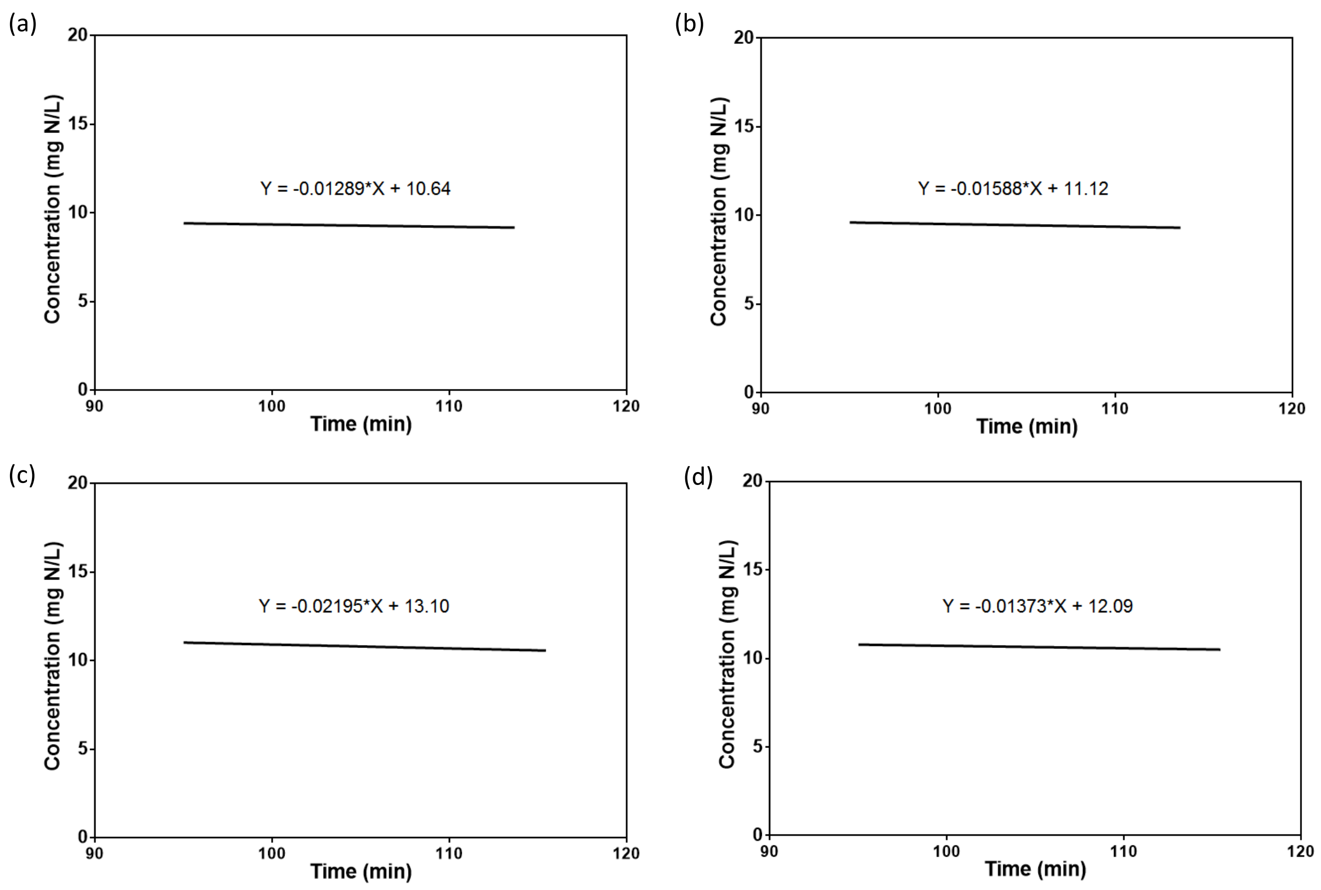


Fig S1. Nitrous oxide ($N_2O$) loss due to stripping or other abiotic phenomena was estimated in control tests (referred to as $r_e$ in Equation 1, Appendix 2). Each control test was conducted in duplicate prior to the batch tests. Panels a,b ,c and d depict the observed $N_2O$ concentration from one of the duplicates for each test.

### Appendix 3

#### 1. *Observed $NO_x$ reduction rates*

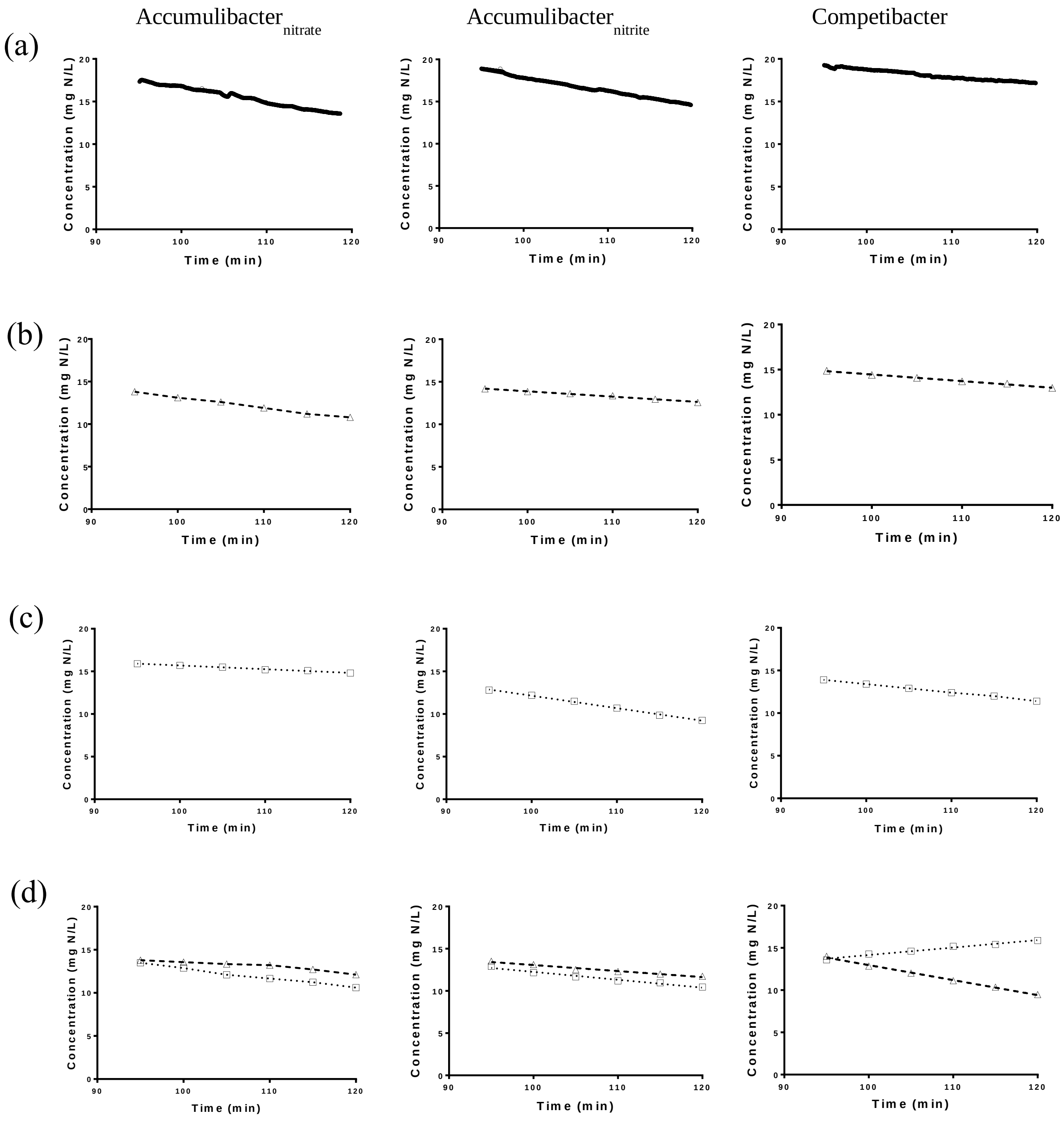

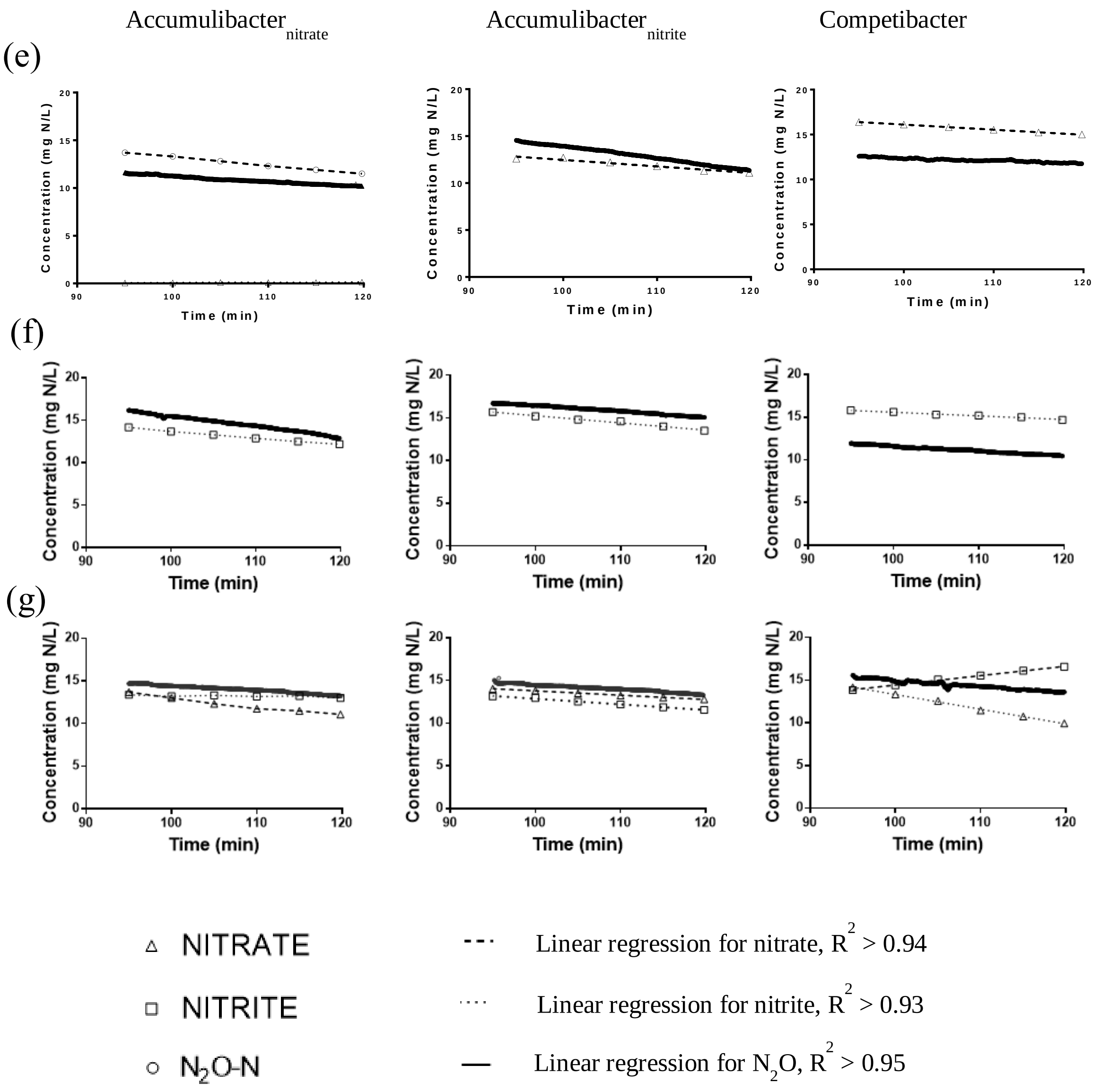


Figure S2. Nitrate ($NO_3^-$), nitrite ($NO_2^-$) and nitrous oxide ($N_2O$) reduction observed for one set of batch experiments with Accumulibacter$_{nitrate}$, Accumulibacter$_{nitrite}$ and Competibacter. The tests shown in the figure are (a) Test A, $N_2O$ only, (b) Test B, $NO_3^-$ only, (c) Test C, $NO_2^-$ only, (d) Test D, $NO_3^-$ and $NO_2^-$, (e) Test E, $NO_3^-$ and $N_2O$, (f) Test F, $NO_2^-$ and $N_2O$, (g) Test G, All.

*2. Electron distribution profile*

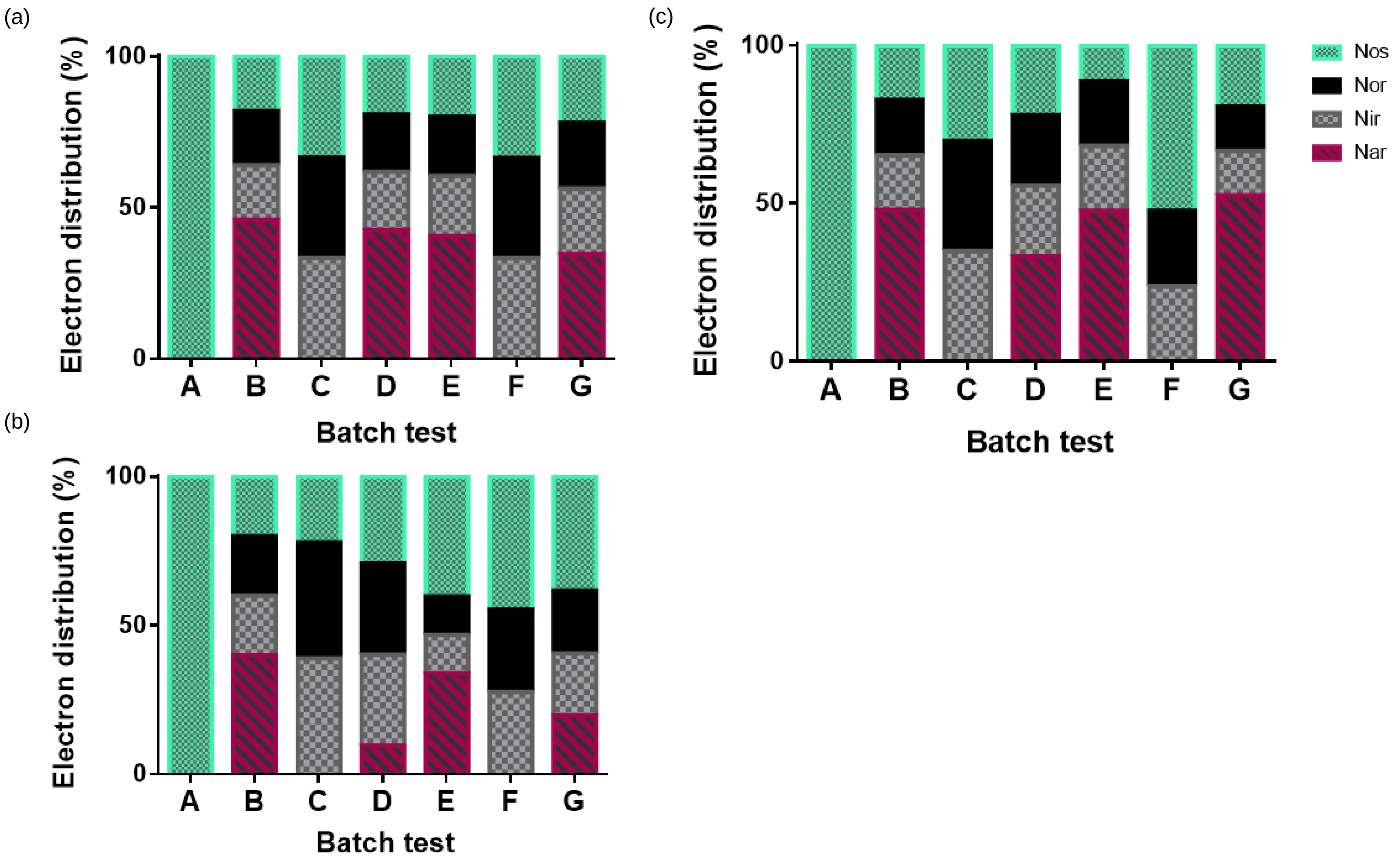

Figure S3. Electron distribution (%) by specific denitrification enzymes for batch test conditions A to G with (a) $Accumulibacter_{nitrite}$ (b) $Accumulibacter_{nitrate}$ (c) Competibacter (d) Defluviicoccus. The different denitrification enzymes are nitrate reductase (Nar, shown in maroon), nitrite reductase (Nir, shown in grey), nitric oxide reductase (Nor, shown in black) and nitrous oxide reductase (Nos, shown in green).

Table S1. Kinetic and stoichiometric parameters for DPAO cultures grown with $NO_3^-$ and $NO_2^-$.

| Parameters | Definition | Values ($NO_3^-$) | Values ($NO_2^-$) | Unit | Source |
|---|---|---|---|---|---|
| Kinetic parameters | | | | | |
| $q_{PHA}$ | Rate constant for storage of $X_{PHA}$ | 0.53 | 0.53 | $h^{-1}$ | [5] |
| $K_{S,DPAO}$ | Biomass saturation constant for $S_S$ | 10 | 10 | mg COD $L^{-3}$ | [6] |
| $K_{PP,DPAO}$ | Saturation constant for $X_{PP}/X_{DPAO}$ | 0.05 | 0.05 | mg P $mg^{-1}$ COD | [6] |
| $q_{PP}$ | Rate constant for storage of $X_{PP}$ | 0.135 ± 0.046 | 0.112 ± 0.022 | $h^{-1}$ | This study |
| $K_{PO_4^{3-},PP}$ | Saturation constant for $S_{PO_4}3-$ of storage | 0.2 | 0.2 | mg P $L^{-3}$ | [6] |
| $K_{PHA}$ | Saturation constant for $X_{PHA}/X_{DPAO}$ | 0.1 | 0.1 | mg COD $mg^{-1}$ COD | [6] |
| $K_{max,DPAO}$ | Maximum ratio of $X_{PP}/X_{DPAO}$ | 0.2 | 0.2 | mg P $mg^{-1}$ COD | [6] |
| $K_{iPP,DPAO}$ | Saturation constant for $K_{max,DPAO}$ - $X_{PP}/X_{DPAO}$ | 0.05 | 0.05 | mg P $mg^{-1}$ COD | [6] |
| $K_{DPAO,PO_4^{3-}}$ | Saturation constant for $S_{PO_4}3-$ of growth | 0.05 | 0.05 | mg P $L^{-3}$ | [6] |
| $\mu_{DPAO1}$ | Anoxic growth rate on nitrate | 0.029 ± 0.007 | 0.035 ± 0.01 | $h^{-1}$ | This study |
| $\mu_{DPAO2}$ | Anoxic growth rate on nitrite | 0.021 ± 0.009 | 0.036 ± 0.006 | $h^{-1}$ | This study |
| $\mu_{DPAO3}$ | Anoxic growth rate on NO | 0.142 | 0.142 | $h^{-1}$ | [7] |
| $\mu_{DPAO4}$ | Anoxic growth rate for on $N_2O$ | 0.066 ± 0.019 | 0.068 ± 0.027 | $h^{-1}$ | This study |
| $K_{NO_3^-}$ | $S_{NO_3}$- affinity constant | 0.251 | 0.251 | mg N $L^{-3}$ | [7] |
| $K_{NO_2^-}$ | $S_{NO_2}$- affinity constant | 0.10 | 0.10 | mg N $L^{-3}$ | This study |
| $K_{NO}$ | $S_{NO}$ affinity constant | 0.0021 | 0.0021 | mg N $L^{-3}$ | [7] |
| $K_{N_2O}$ | $S_{N_2O}$ affinity constant | 0.0052 | 0.0052 | mg N $L^{-3}$ | [7] |
| $b_{DPAO}$ | Endogenous respiration rate of $X_{DPAO}$ | 0.005 | 0.005 | $h^{-1}$ | [6] |
| $b_{PP}$ | Lysis of $X_{PP}$ | 0.005 | 0.005 | $h^{-1}$ | [6] |
| $b_{PHA}$ | Respiration rate for $X_{PHA}$ | 0.005 | 0.005 | $h^{-1}$ | [6] |
| $K_{NO_x}$ | Biomass $S_{NO_x}$ affinity | 0.5 | 0.5 | mg N $L^{-3}$ | [6] |

| | | | | | |
|---|---|---|---|---|---|
| | constant | | | | |
| Stoichiometric parameters | | | | | |
| $Y_{PO_4^{3-}}$ | Requirement of $X_{PP}$ per XPHA storage | 0.3 | 0.3 | mg P mg$^{-1}$ COD | [6] |
| $Y_{PHA}$ | Requirement of $X_{PHA}$ per $X_{PP}$ storage | 0.2 | 0.2 | mg COD mg$^{-1}$ P | [6] |
| $Y_{DPAO,NO_x}$ | Anoxic yield coefficient for $X_{DPAO}$ | 0.5 | 0.5 | mg COD mg$^{-1}$ COD | [6] |
| $i_{P,BM}$ | Phosphorus content of biomass | 0.02 | 0.02 | mg P mg$^{-1}$ COD | [6] |
| $i_{P,XI}$ | Phosphorus content of $X_I$ | 0.01 | 0.01 | mg P mg$^{-1}$ COD | [6] |
| $f_I$ | Fraction of $X_I$ in endogenous respiration | 0.2 | 0.2 | mg COD mg$^{-1}$ COD | [6] |

Table S2. Kinetic rate expressions for the DPAO model (as per Liu *et al.* 2015).

| Process | Rate expressions |
|---|---|
| 1. Anaerobic storage of $X_{PHA}$ | $q_{PHA}\frac{S_S}{K_{S,DPAO}+S_S}\frac{X_{PP}/X_{DPAO}}{K_{PP,DAPO}+X_{PP}/X_{DPAO}}X_{DPAO}$ |
| 2. Anoxic storage of $X_{PP}$ on $NO_3^-$ | $q_{PP}\frac{S_{NO_3^-}}{K_{NO_3^-}+S_{NO_3^-}}\frac{S_{PO_4^{3-}}}{K_{PO_4^{3-},PP}+S_{PO_4^{3-}}}\frac{X_{PHA}/X_{DPAO}}{K_{PHA}+X_{PHA}/X_{DPAO}}\frac{K_{max,DPAO}-X_{PP}/X_{DPAO}}{K_{iPP,DPAO}+K_{max,DPAO}-X_{PP}/X_{DPAO}}X_{DPAO}$ |
| 3. Anoxic storage of $X_{PP}$ on $NO_2^-$ | $q_{PP}\frac{S_{NO_2^-}}{K_{NO_2^-}+S_{NO_2^-}}\frac{S_{PO_4^{3-}}}{K_{PO_4^{3-},PP}+S_{PO_4^{3-}}}\frac{X_{PHA}/X_{DPAO}}{K_{PHA}+X_{PHA}/X_{DPAO}}\frac{K_{max,DPAO}-X_{PP}/X_{DPAO}}{K_{iPP,DPAO}+K_{max,DPAO}-X_{PP}/X_{DPAO}}X_{DPAO}$ |
| 4. Anoxic storage of $X_{PP}$ on NO | $q_{PP}\frac{S_{NO}}{K_{NO}+S_{NO}}\frac{S_{PO_4^{3-}}}{K_{PO_4^{3-},PP}+S_{PO_4^{3-}}}\frac{X_{PHA}/X_{DPAO}}{K_{PHA}+X_{PHA}/X_{DPAO}}\frac{K_{max,DPAO}-X_{PP}/X_{DPAO}}{K_{iPP,DPAO}+K_{max,DPAO}-X_{PP}/X_{DPAO}}X_{DPAO}$ |
| 5. Anoxic storage of $X_{PP}$ on $N_2O$ | $q_{PP}\frac{S_{N_2O}}{K_{N_2O}+S_{N_2O}}\frac{S_{PO_4^{3-}}}{K_{PO_4^{3-},PP}+S_{PO_4^{3-}}}\frac{X_{PHA}/X_{DPAO}}{K_{PHA}+X_{PHA}/X_{DPAO}}\frac{K_{max,DPAO}-X_{PP}/X_{DPAO}}{K_{iPP,DPAO}+K_{max,DPAO}-X_{PP}/X_{DPAO}}X_{DPAO}$ |
| 6. Anoxic growth on $NO_3^-$ | $\mu_{DPAO1}\frac{S_{NO_3^-}}{K_{NO_3^-}+S_{NO_3^-}}\frac{S_{PO_4^{3-}}}{K_{DPAO,PO_4^{3-}}+S_{PO_4^{3-}}}\frac{X_{PHA}/X_{DPAO}}{K_{PHA}+X_{PHA}/X_{DPAO}}X_{DPAO}$ |
| 7. Anoxic growth on $NO_2^-$ | $\mu_{DPAO2}\frac{S_{NO_2^-}}{K_{NO_2^-}+S_{NO_2^-}}\frac{S_{PO_4^{3-}}}{K_{DPAO,PO_4^{3-}}+S_{PO_4^{3-}}}\frac{X_{PHA}/X_{DPAO}}{K_{PHA}+X_{PHA}/X_{DPAO}}X_{DPAO}$ |
| 8. Anoxic growth on NO | $\mu_{DPAO3}\frac{S_{NO}}{K_{NO}+S_{NO}}\frac{S_{PO_4^{3-}}}{K_{DPAO,PO_4^{3-}}+S_{PO_4^{3-}}}\frac{X_{PHA}/X_{DPAO}}{K_{PHA}+X_{PHA}/X_{DPAO}}X_{DPAO}$ |
| 9. Anoxic growth on $N_2O$ | $\mu_{DPAO4}\frac{S_{N_2O}}{K_{N_2O}+S_{N_2O}}\frac{S_{PO_4^{3-}}}{K_{DPAO,PO_4^{3-}}+S_{PO_4^{3-}}}\frac{X_{PHA}/X_{DPAO}}{K_{PHA}+X_{PHA}/X_{DPAO}}X_{DPAO}$ |
| 10. Anoxic endogenous respiration on $NO_3^-$ | $b_{DPAO}\frac{S_{NO_3^-}}{K_{NO_x}+S_{NO_3^-}}X_{DPAO}$ |

11. Anoxic endogenous respiration on $NO_2^-$ $$b_{DPAO}\frac{S_{NO_2^-}}{K_{NO_x}+S_{NO_2^-}}X_{DPAO}$$
12. Anoxic endogenous respiration on NO $$b_{DPAO}\frac{S_{NO}}{K_{NO_x}+S_{NO}}X_{DPAO}$$
13. Anoxic endogenous respiration on $N_2O$ $$b_{DPAO}\frac{S_{N_2O}}{K_{NO_x}+S_{N_2O}}X_{DPAO}$$
14. Anoxic respiration of $X_{PP}$ on $NO_3^-$ $$b_{PP}\frac{S_{NO_3^-}}{K_{NO_x}+S_{NO_3^-}}X_{PP}$$
15. Anoxic respiration of $X_{PP}$ on $NO_2^-$ $$b_{PP}\frac{S_{NO_2^-}}{K_{NO_x}+S_{NO_2^-}}X_{PP}$$
16. Anoxic respiration of $X_{PP}$ on NO $$b_{PP}\frac{S_{NO}}{K_{NO_x}+S_{NO}}X_{PP}$$
17. Anoxic respiration of $X_{PP}$ on $N_2O$ $$b_{PP}\frac{S_{N_2O}}{K_{NO_x}+S_{N_2O}}X_{PP}$$
18. Anoxic respiration of $X_{PHA}$ on $NO_3^-$ $$b_{PHA}\frac{S_{NO_3^-}}{K_{NO_x}+S_{NO_3^-}}X_{PHA}$$
19. Anoxic respiration of $X_{PHA}$ on $NO_2^-$ $$b_{PHA}\frac{S_{NO_2^-}}{K_{NO_x}+S_{NO_2^-}}X_{PHA}$$
20. Anoxic respiration of $X_{PHA}$ on NO $$b_{PHA}\frac{S_{NO}}{K_{NO_x}+S_{NO}}X_{PHA}$$
21. Anoxic respiration of $X_{PHA}$ on $N_2O$ $$b_{PHA}\frac{S_{N_2O}}{K_{NO_x}+S_{N_2O}}X_{PHA}$$

Table S3. Peterson matrix for the DPAO model (as per Liu *et al.* 2015).

| Variable | $S_{NO_3^-}$ | $S_{NO_2^-}$ | $S_{NO}$ | $S_{N_2O}$ | $S_{N_2}$ | $S_S$ | $S_{PO_4^{3-}}$ | $X_{DPAO}$ | $X_{PP}$ | $X_{PHA}$ | $X_I$ |
|---|---|---|---|---|---|---|---|---|---|---|---|
| Process | N | N | N | N | N | COD | P | COD | P | COD | COD |
| 1. Anaerobic storage of $X_{PHA}$ | | | | | | -1 | $Y_{PO_4^{3-}}$ | | $-Y_{PO_4^{3-}}$ | 1 | |
| 2. Anoxic storage of $X_{PP}$ on $NO_3^-$ | $-\frac{Y_{PHA}}{1.14}$ | $\frac{Y_{PHA}}{1.14}$ | | | | | -1 | | 1 | $-Y_{PHA}$ | |
| 3. Anoxic storage of $X_{PP}$ on $NO_2^-$ | | $-\frac{Y_{PHA}}{0.57}$ | $\frac{Y_{PHA}}{0.57}$ | | | | -1 | | 1 | $-Y_{PHA}$ | |
| 4. Anoxic storage of $X_{PP}$ on NO | | | $-\frac{Y_{PHA}}{0.57}$ | $\frac{Y_{PHA}}{0.57}$ | | | -1 | | 1 | $-Y_{PHA}$ | |
| 5. Anoxic storage of $X_{PP}$ on $N_2O$ | | | | | $-\frac{Y_{PHA}}{0.57}$ | $\frac{Y_{PHA}}{0.57}$ | -1 | | 1 | $-Y_{PHA}$ | |
| 6. Anoxic growth on $NO_3^-$ | $-\frac{1-Y_{DPA}}{1.14Y_{DPA}}$ | $\frac{1-Y_{DPAO,N}}{1.14Y_{DPAO,N}}$ | | | | | $-i_{P,BM}$ | 1 | | $-\frac{1}{Y_{DPAO,NO_X}}$ | |
| 7. Anoxic growth on $NO_2^-$ | | $-\frac{1-Y_{DPAO}}{0.57Y_{DPAO}}$ | $\frac{1-Y_{DPAO,N}}{0.57Y_{DPAO,N}}$ | | | | $-i_{P,BM}$ | 1 | | $-\frac{1}{Y_{DPAO,NO_X}}$ | |
| 8. Anoxic growth on NO | | | $-\frac{1-Y_{DPAO,}}{0.57Y_{DPAO}}$ | $\frac{1-Y_{DPAO,NO_X}}{0.57Y_{DPAO,NO_X}}$ | | | $-i_{P,BM}$ | 1 | | $-\frac{1}{Y_{DPAO,NO_X}}$ | |
| 9. Anoxic growth on $N_2O$ | | | | $-\frac{1-Y_{DPAO,N}}{0.57Y_{DPAO,N}}$ | $\frac{1-Y_{DPAO,N}}{0.57Y_{DPAO,N}}$ | | $-i_{P,BM}$ | 1 | | $-\frac{1}{Y_{DPAO,NO_X}}$ | |
| 10. Anoxic endogenous respiration on $NO_3^-$ | $-\frac{1-f_I}{1.14}$ | $\frac{1-f_I}{1.14}$ | | | | | $i_{P,BM} - i_{P,XI}f_i$ | -1 | | | $f_I$ |
| 11. Anoxic endogenous respiration on | | $-\frac{1-f_I}{0.57}$ | $\frac{1-f_I}{0.57}$ | | | | $i_{P,BM} - i_{P,XI}f_i$ | -1 | | | $f_I$ |

| | | | | | | | | | | |
|---|---|---|---|---|---|---|---|---|---|---|
| $NO_2^-$ | | | | | | | | | | |
| 12. Anoxic endogenous respiration on NO | | | $-\frac{1-f_I}{0.57}$ | $\frac{1-f_I}{0.57}$ | | $i_{P,BM} - i_{P,XI}f_i$ | -1 | | | $f_I$ |
| 13. Anoxic endogenous respiration on $N_2O$ | | | | $-\frac{1-f_I}{0.57}$ | $\frac{1-f_I}{0.57}$ | $i_{P,BM} - i_{P,XI}f_i$ | -1 | | | $f_I$ |
| 14. Anoxic respiration of $X_{PP}$ on $NO_3^-$ | | | | | | 1 | | -1 | | |
| 15. Anoxic respiration of $X_{PP}$ on $NO_2^-$ | | | | | | 1 | | -1 | | |
| 16. Anoxic respiration of $X_{PP}$ on NO | | | | | | 1 | | -1 | | |
| 17. Anoxic respiration of $X_{PP}$ on $N_2O$ | | | | | | 1 | | -1 | | |
| 18. Anoxic respiration of $X_{PHA}$ on $NO_3^-$ | $-\frac{1}{1.14}$ | $\frac{1}{1.14}$ | | | | | | | -1 | |
| 19. Anoxic respiration of $X_{PHA}$ on $NO_2^-$ | | $-\frac{1}{0.57}$ | $\frac{1}{0.57}$ | | | | | | -1 | |
| 20. Anoxic respiration of $X_{PHA}$ on NO | | | $-\frac{1}{0.57}$ | $\frac{1}{0.57}$ | | | | | -1 | |
| 21. Anoxic respiration of $X_{PHA}$ on $N_2O$ | | | | $-\frac{1}{0.57}$ | $\frac{1}{0.57}$ | | | | -1 | |

Table S4. Model fitting parameters for $Accumulibacter_{nitrate}$

| | $NO_3^-$ | | | $NO_2^-$ | | | $N_2O$ | | | $PO_4^{3-}$ | | |
|---|---|---|---|---|---|---|---|---|---|---|---|---|
| Electron acceptor | $R^2$ | $R_{Adj}^2$ | RMSE | $R^2$ | $R_{Adj}^2$ | RMSE | $R^2$ | $R_{Adj}^2$ | RMSE | $R^2$ | $R_{Adj}^2$ | RMSE |
| $NO_3^-$ | 0.998 | 0.997 | 0.0332 | 0.938 | 0.922 | 0.039 | | | | 0.995 | 0.994 | 0.133 |
| $NO_2^-$ | | | | 0.989 | 0.987 | 0.087 | 0.937 | 0.922 | 0.014 | 0.978 | 0.973 | 0.214 |
| $N_2O$ | | | | | | | 0.995 | 0.994 | 0.125 | 0.988 | 0.985 | 0.158 |
| $NO_3^-$ + $NO_2^-$ | 0.995 | 0.994 | 0.063 | 0.994 | 0.993 | 0.056 | 0.858 | 0.822 | 0.020 | 0.998 | 0.997 | 0.063 |
| $NO_3^-$ + $N_2O$ | 0.998 | 0.997 | 0.0415 | 0.847 | 0.809 | 0.025 | 0.985 | 0.982 | 0.105 | 0.995 | 0.993 | 0.103 |
| $NO_2^-$ + $N_2O$ | | | | 0.996 | 0.996 | 0.051 | 0.977 | 0.972 | 0.190 | 0.966 | 0.957 | 0.252 |
| $NO_3^-$ + $NO_2^-$ + $N_2O$ | 0.983 | 0.979 | 0.104 | 0.996 | 0.995 | 0.042 | 0.977 | 0.971 | 0.098 | 0.92 | 0.9 | 0.357 |

Table S5. Model fitting parameters for $Accumulibacter_{nitrite}$

| | $NO_3^-$ | | | $NO_2^-$ | | | $N_2O$ | | | $PO_4^{3-}$ | | |
|---|---|---|---|---|---|---|---|---|---|---|---|---|
| Electron acceptor | $R^2$ | $R_{Adj}^2$ | RMSE | $R^2$ | $R_{Adj}^2$ | RMSE | $R^2$ | $R_{Adj}^2$ | RMSE | $R^2$ | $R_{Adj}^2$ | RMSE |
| $NO_3^-$ | 0.885 | 0.857 | 0.226 | | | | | | | 0.973 | 0.966 | 0.134 |
| $NO_2^-$ | | | | 0.989 | 0.987 | 0.138 | 0.988 | 0.985 | 0.049 | 0.928 | 0.91 | 0.333 |
| $N_2O$ | | | | | | | 0.999 | 0.999 | 0.047 | 0.99 | 0.987 | 0.122 |
| $NO_3^-$ + $NO_2^-$ | 0.991 | 0.989 | 0.0655 | 0.969 | 0.961 | 0.133 | 1.000 | 0.990 | 0.004 | 0.88 | 0.85 | 0.493 |
| $NO_3^-$ + $N_2O$ | 0.934 | 0.918 | 0.203 | 0.899 | 0.874 | 0.060 | 0.961 | 0.952 | 0.223 | 0.948 | 0.935 | 0.196 |
| $NO_2^-$ + $N_2O$ | | | | 0.996 | 0.995 | 0.075 | 0.962 | 0.952 | 0.160 | 0.942 | 0.928 | 0.463 |
| $NO_3^-$ + $NO_2^-$ + $N_2O$ | 0.973 | 0.966 | 0.123 | 0.967 | 0.959 | 0.144 | 0.983 | 0.979 | 0.112 | 0.875 | 0.844 | 0.734 |

**Table S6.** Best fit parameters for anoxic growth rate for each denitrification step with 95% confidence intervals describing the multi-step denitrification process.

| **Parameters** | **Literature values** | **Measured anoxic growth rate ($h^{-1}$) in batch test** | | | | | | |
|---|---|---|---|---|---|---|---|---|
| | | Test A ($N_2O$) | Test B ($NO_3^-$) | Test C ($NO_2^-$) | Test D ($NO_2^-$& $NO_3^-$) | Test E ($NO_3^-$& $N_2O$) | Test F ($NO_2^-$& $N_2O$) | Test G ($NO_2^-$, $NO_3^-$ & $N_2O$) |
| **Accumulibacter$_{nitrite}$** | | | | | | | | |
| **m_DPAO1** | 0.07 | - | 0.02 | - | 0.03 | 0.04 | - | 0.028 |
| **m_DPAO2** | 0.019 | - | 0.018 | 0.015 | 0.025 | 0.0375 | 0.0075 | 0.025 |
| **m_DPAO3** | 0.142 | - | 0.14 | 0.14 | 0.142 | 0.142 | 0.142 | 0.142 |
| **m_DPAO4** | 0.018 | 0.07 | 0.05 | 0.05 | 0.09 | 0.04 | 0.09 | 0.075 |
| **Accumulibacter$_{nitrate}$** | | | | | | | | |
| **m_DPAO1** | 0.07 | - | 0.025 | - | 0.0375 | 0.05 | - | 0.028 |
| **m_DPAO2** | 0.019 | - | 0.025 | 0.04 | 0.045 | 0.035 | 0.0275 | 0.03 |
| **m_DPAO3** | 0.142 | - | 0.142 | 0.142 | 0.142 | 0.142 | 0.142 | 0.142 |
| **m_DPAO4** | 0.018 | 0.02 | 0.025 | 0.04 | 0.08 | 0.09 | 0.085 | 0.09 |
